# Predictive pre-activation of orthographic and lexical-semantic representations facilitates visual word recognition

**DOI:** 10.1101/2020.07.14.202226

**Authors:** Susanne Eisenhauer, Benjamin Gagl, Christian J. Fiebach

**Affiliations:** Department of Psychology, Goethe University Frankfurt, Frankfurt am Main, Germany; Center for Individual Development and Adaptive Education of Children at Risk (IDeA), Frankfurt am Main, Germany; Brain Imaging Center, Goethe University Frankfurt, Frankfurt am Main, Germany

## Abstract

To a crucial extent, the efficiency of reading results from the fact that visual word recognition is faster in predictive contexts. Predictive coding models suggest that this facilitation results from pre-activation of predictable stimulus features across multiple representational levels before stimulus onset. Still, it is not sufficiently understood which aspects of the rich set of linguistic representations that are activated during reading – visual, orthographic, phonological, and/or lexical-semantic – contribute to context-dependent facilitation. To investigate in detail which linguistic representations are pre-activated in a predictive context and how they affect subsequent stimulus processing, we combined a well-controlled repetition priming paradigm, including words and pseudowords (i.e., pronounceable nonwords), with behavioral and magnetoencephalography measurements. For statistical analysis, we used linear mixed modeling, which we found had a higher statistical power compared to conventional multivariate pattern decoding analysis. Behavioral data from 49 participants indicate that word predictability (i.e., context present vs. absent) facilitated orthographic and lexical-semantic, but not visual or phonological processes. Magnetoencephalography data from 38 participants show sustained activation of orthographic and lexical-semantic representations in the interval before processing the predicted stimulus, suggesting selective pre-activation at multiple levels of linguistic representation as proposed by predictive coding. However, we found more robust lexical-semantic representations when processing predictable in contrast to unpredictable letter strings, and pre-activation effects mainly resembled brain responses elicited when processing the expected letter string. This finding suggests that pre-activation did not result in ‘explaining away’ predictable stimulus features, but rather in a ‘sharpening’ of brain responses involved in word processing.

## 1 Introduction

Predictive contexts can facilitate word processing, in the sense of increasing reading speed (e.g., Rayner et al., 2011) and decreasing neuronal activation elicited during recognition of expected words (as reflected, for example, in the reduction of the N400 component of the event-related brain potential; e.g., Kutas and Hillyard, 1984; Kutas and Federmeier, 2011). These often-replicated findings have been linked to domain-general theories that postulate active, top-down (i.e., from hierarchically higher to lower processing levels) prediction of expected stimulus characteristics before actually perceiving the stimulus (predictive coding; cf. Rao and Ballard, 1999; Friston, 2005). In line with this, several neuro-cognitive models of visual word recognition (e.g., Seidenberg and McClelland, 1989; Carreiras et al., 2014) assume context-based prediction across multiple levels of linguistic processing, and it has been hypothesized that hierarchical predictions during reading involve the pre-activation of visual, pre-lexical (i.e., orthographic or phonological), and lexical-semantic representations of predicted words (Federmeier, 2007; Kuperberg and Jaeger, 2016). However, this proposal has not been systematically tested because currently, available evidence does not unambiguously differentiate between predictive pre-activation of representations at these different linguistic processing stages.

First studies showed that language-related brain regions can be activated before a highly expected stimulus appears (Dikker and Pylkkanen, 2011; Bonhage et al., 2015; Wang et al., 2017), but did not further assess the nature of pre-activated representations. More recent work showed pre-stimulus effects of semantic category (Heikel et al., 2018; Wang et al., 2020) using multivariate pattern analysis techniques (King et al., 2018; Kragel et al., 2018), as well as pre-stimulus effects of word frequency (Fruchter et al., 2015) using linear mixed models (LMMs; Baayen et al., 2008). These studies provide initial evidence that lexical-semantic representations are pre-activated before a predictable word appears. Also, several studies manipulating lexical-semantic context (e.g., the sentence context preceding the target word) found context-based modulations of brain activation in time windows associated with visual or pre-lexical processing (e.g., Lee et al., 2012; Brothers et al., 2015), which however provides only indirect evidence for predictive processing at other levels of linguistic representation than the lexical-semantic level. Studies that investigated pre-lexical context effects more directly found no (Eisenhauer et al., 2019) or only minimal effects (Nieuwland et al., 2018; Nicenboim et al., 2020), indicating that more statistically robust approaches may be needed for identifying the processes involved in pre-lexical pre-activation (see also Nieuwland, 2019, for a review).

Here, we combine behavioral and magnetoencephalography (MEG) data elicited during processing of words and (orthographically legal and pronounceable) pseudowords, to investigate the mechanisms of predictive pre-activation at multiple levels of linguistic representation, i.e., visual, pre-lexical (orthographic and phonological), and lexical-semantic. Contextual predictability was explicitly controlled by using a repetition priming paradigm, which is a common approach for investigating predictive processing (e.g., Auksztulewicz and Friston, 2016; Grotheer and Kovács, 2016). In a first step, we conducted a behavioral experiment to determine which representational levels of visual word recognition are influenced by predictive processing. In detail, we determined whether context-dependent facilitation (‘priming’) interacts with quantitative metrics from psycholinguistics representing different stages of word processing, i.e., visual stimulus complexity (early visual processing; e.g., Pelli et al., 2006), orthographic word similarity (pre-lexical orthographic processing; e.g., Yarkoni et al., 2008a), the number of syllables (pre-lexical phonological processing; e.g., Álvarez et al., 2010), and word frequency and stimulus lexicality (lexical-semantic processing; e.g., Forster and Chambers, 1973; Fiebach et al., 2002). Subsequently, MEG activity measured in an independent experiment was explored strictly for those metrics that interacted in behavior with contextual predictability (i.e., whose effects differed between primed vs. unprimed words). This procedure prevented the investigation of neurophysiological effects without a behavioral counterpart, which would be difficult to interpret (Krakauer et al., 2017).

We investigated MEG data given its excellent temporal resolution (Gross, 2019) to separate effects of predictive pre-activation and stimulus processing. We first investigated the neurophysiological correlates of predictability across representational levels *during* stimulus processing by assessing the interaction of context effects with psycholinguistic metrics. Based on the observed pattern of context effects, we were able to differentiate whether predictive processing in visual word recognition is based on predictive coding (according to which predictable stimulus features are ‘explained away’) or on a ‘sharpening’ mechanism (which postulates the suppression of noise for predictable stimuli; cf. Kok et al., 2012; Blank and Davis, 2016). Crucially, we also assessed effects of psycholinguistic metrics in the delay period *prior to* predictable target letter strings. Detecting these effects would provide direct evidence for predictive pre-activation at the associated representational level. As a critical test case for a mechanistic contribution of the respective linguistic processing stage to context-dependent predictability, we hypothesized that the strength of pre-activation effects in the delay should be inversely related to the strength of neural effects measured during processing of the target. Finally, we localized the brain regions underlying the observed effects to assess whether the brain regions implicated in letter string processing are also implicated in predictive pre-activation.

## 2 Methods

### 2.1 Behavioral experiment

#### 2.1.1 Participants

Forty-nine healthy, right-handed, native speakers of German recruited from university campuses (33 females, mean age 24.7 ± 4.9 years, range: 18–39 years) were included in the final data analyses. All participants had normal or corrected-to-normal vision, and normal reading abilities as assessed with the adult version of the Salzburg Reading Screening (the unpublished adult version of Mayringer and Wimmer, 2003). Further participants were excluded before the experiment due to low reading skills (i.e., reading test score below 16th percentile; N = 3) or participation in a similar previous experiment (N = 1), and during the course of the experiment due to failure to complete the experimental protocol (N = 8) and because of an experimenter error (mix-up of the pseudoword lists during the pre-experiment familiarization procedure; N = 1). All participants gave written informed consent according to procedures approved by the local ethics committee (Department of Psychology, Goethe University Frankfurt, application N° 2015-229) and received 10 € per hour or course credit as compensation.

#### 2.1.2 Stimuli and presentation procedure

60 words and 180 pseudowords (five letters each) were presented (black on white background, 14 pt., 51 cm viewing distance) in a repetition priming experiment consisting of two priming blocks and two non-priming blocks (120 trials per block; Fig. 1a) with a total duration of ∼20 min. This paradigm allows for strong predictions while maintaining the aspect of natural reading that letter strings are processed sequentially. For the present study, we focused on a subset of 60 pseudowords that were unfamiliar to the participants (‘novel pseudoword’ condition). However, note that around 90 min prior to the priming experiment described here, the pseudowords had been presented to the participants in another experiment not of interest for the present study, without any learning instruction. In addition, the experiment contained two further sets of 60 pseudowords each, which were familiarized prior to the experiment as described in Eisenhauer et al. (2019; behavioral experiment). A description of the learning procedure and the results can be found in Supporting Results 2. Participants learned meanings for one of these pseudoword sets (i.e., ‘semantic pseudowords’); however, this set was not part of the present analysis as no comparable condition was included in the MEG dataset. The pseudowords from the final set were familiarized via repeated presentation and reading aloud without learning a meaning for these pseudowords. Thus, these ‘familiar pseudowords’ were not associated with lexical-semantic representations, but were nevertheless familiarized at a pre-lexical level as participants gained familiarity with the orthographic and phonological structure of these pseudowords. These pseudowords were included in a control analysis (see below). The assignment of pseudoword sets to the three conditions (i.e., novel pseudowords, familiar pseudowords and semantic pseudowords) was varied across participants.

**Figure 1.**
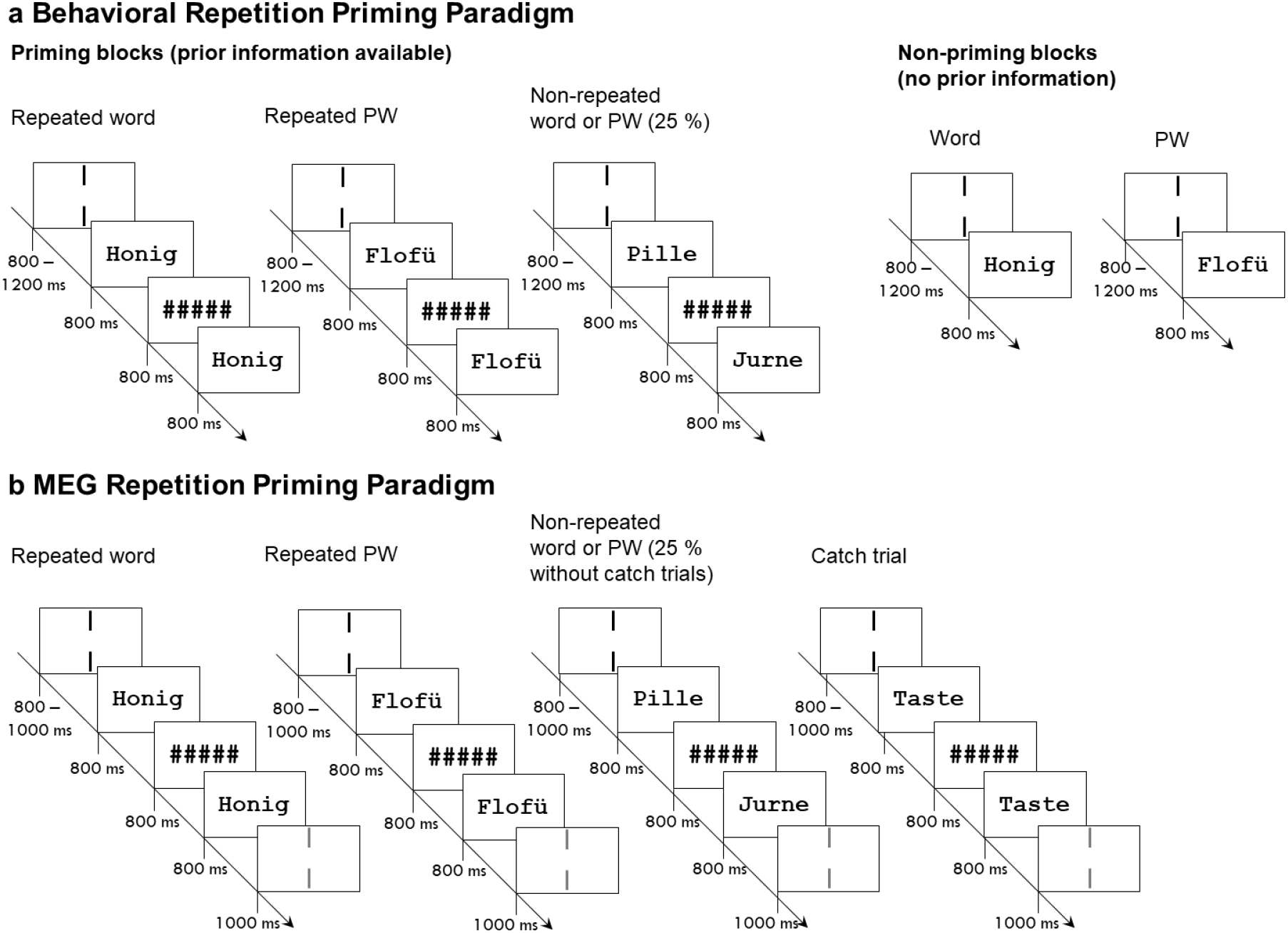
Experimental procedures. (a) Behavioral repetition priming paradigm. Two priming blocks (left) and two non-priming blocks (right) were presented in alternating order. In priming blocks, each trial consisted of a prime and a target stimulus presented for 800 ms each, separated by an interval of 800 ms during which a string of five hash marks was presented. Stimuli could be words or pseudowords (PW). 75 % of trials were repetition trials with identical prime and target, while in the remaining 25 % two different letter strings were presented (non-repetition trials; not analyzed). In this case, prime and target could be from the same or from two different conditions, with all combinations of conditions appearing equally often. In non-priming blocks, only one word or pseudoword was presented in each trial. Participants were instructed to respond on each trial whether or not they had a semantic association with the target in priming blocks or with the isolated item in non-priming blocks. Before onset of the prime or the isolated item, two black vertical bars presented for 800 – 1,200 ms indicated the center of the screen where participants were asked to fixate. Context effects were investigated by comparing isolated items from the non-priming blocks with the targets from the repetition trials. (b) Repetition priming paradigm during MEG recording. The presentation procedure was identical to the priming blocks in (a). There were no non-priming blocks. Additionally, after target offset two grey vertical bars were presented for 1,000 ms indicating a blinking period. Before the onset of the next trial, a blank screen was presented for 500 ms. Participants were instructed to silently read presented letter strings and to respond only to rare catch trials (i.e., presentation of the word Taste, Engl. button). Context effects were investigated by comparing primes to repeated targets.

In priming blocks, a prime and a target stimulus were presented for 800 ms each, separated by a delay period of 800 ms during which a string of five hash marks was presented. Prime and target were identical in 75 % of trials. In non-priming blocks, a single letter string was presented for 800 ms in each trial. We choose extended presentation durations for the MEG study to separate effects of stimulus processing and pre-activation, which we expected during the delay period. Thus, for comparability, we also adopted the timing for the behavioral study. The inter-trial-interval was jittered between 800 and 1,200 ms (mean: 1,000 ms). Participants were asked to fixate the space between two vertical black bars at the center of the screen. Upon presentation of a letter string between the two lines, they had to indicate as quickly and accurately as possible whether it had a semantic association or not (which was the case for 50 % of items, i.e., for words and for one list of pseudowords that had been semantically familiarized prior to the experiment). For simplicity, these judgments will be called lexical decisions in the following. The novel and familiar pseudowords used for the present analyses had no semantic associations. In priming blocks, participants responded only to the second (i.e., the target) letter string. Response hands and the order of blocks were counterbalanced across participants. Each letter string was presented in one priming trial and one non-priming trial.

Words and pseudowords were matched between lists (i) for orthographic similarity (‘word likeness’) using the Orthographic Levenshtein Distance 20 (OLD20; Yarkoni et al., 2008a) based on the SUBTLEX-DE database (Brysbaert et al., 2011) and (ii) with respect to the number of syllables (computed via Balloon, cf. Reichel, 2012; see also Table 1). Other psycholinguistic metrics of interest for our analyses were logarithmic word frequency and trigram frequency (i.e., the mean frequency of each trigram per word; obtained from SUBTLEX-DE), as well as visual complexity measures (perimetric complexity and the number of simple features), which were obtained for each letter from the GraphCom database (Chang et al., 2018) and averaged across the five letters of each stimulus (Table 1). The other two visual complexity parameters from GraphCom, i.e., the number of connected points and the number of disconnected components, were not chosen for our analyses, the former due to its high correlation with the number of simple features (>.8 in our stimuli) and the latter due to its low variance across letters of the German language. For parameter correlations within words and pseudowords, see Figure S1 in Supporting Information.

**Table 1.** Stimulus parameters of words and pseudowords (PW) in the behavioral and MEG experiments

|  | 1<sup>st</sup> |  |  |  |  |  |  |
| --- | --- | --- | --- | --- | --- | --- | --- |
|  | Minimum | Quartile | Median | 3<sup>rd</sup> Quartile | Maximum | Mean | SE |
| **Behavior: Logarithmic word frequency** |  |  |  |  |  |  |  |
| **Words** | 0.000 | 1.518 | 1.971 | 2.189 | 3.301 | 1.933 | 0.095 |
| **Behavior: OLD20** |  |  |  |  |  |  |  |
| **Words** | 1.000 | 1.288 | 1.650 | 1.750 | 1.950 | 1.538 | 0.038 |
| **PW set 1** | 1.000 | 1.500 | 1.650 | 1.762 | 2.000 | 1.605 | 0.032 |
| **PW set 2** | 1.000 | 1.288 | 1.650 | 1.850 | 2.100 | 1.542 | 0.045 |
| **PW set 3** | 1.000 | 1.337 | 1.700 | 1.850 | 2.300 | 1.596 | 0.044 |
| **Behavior: Trigram frequency** |  |  |  |  |  |  |  |
| **Words** | 1.806 | 2.556 | 2.793 | 3.162 | 3.773 | 2.837 | 0.056 |
| **PW set 1** | 1.342 | 2.623 | 2.856 | 3.067 | 3.514 | 2.826 | 0.047 |
| **PW set 2** | 1.176 | 2.567 | 2.833 | 3.136 | 3.136 | 2.792 | 0.063 |
| **PW set 3** | 1.437 | 2.576 | 2.844 | 3.237 | 3.602 | 2.808 | 0.061 |
| **Behavior: Number of syllables** |  |  |  |  |  |  |  |
| **Words** | 1.00 | 2.00 | 2.00 | 2.00 | 3.00 | 1.833 | 0.059 |
| **PW set 1** | 1.00 | 2.00 | 2.00 | 2.00 | 2.00 | 1.95 | 0.028 |
| **PW set 2** | 1.00 | 2.00 | 2.00 | 2.00 | 2.00 | 1.967 | 0.023 |
| **PW set 3** | 1.00 | 2.00 | 2.00 | 2.00 | 2.00 | 1.9 | 0.039 |
| **Behavior: Perimetric complexity** |  |  |  |  |  |  |  |
| **Words** | 4.400 | 5.800 | 6.400 | 6.600 | 8.200 | 6.354 | 0.093 |
| **PW set 1** | 5.000 | 5.800 | 6.300 | 6.800 | 7.800 | 6.320 | 0.083 |
| **PW set 2** | 5.200 | 6.200 | 6.600 | 7.000 | 7.800 | 6.553 | 0.083 |
| **PW set 3** | 5.000 | 5.950 | 6.600 | 7.000 | 7.800 | 6.457 | 0.090 |
| **Behavior: Number of simple features** |  |  |  |  |  |  |  |
| **Words** | 1.400 | 1.950 | 2.000 | 2.200 | 2.600 | 2.031 | 0.031 |
| **PW set 1** | 1.200 | 1.800 | 2.000 | 2.200 | 2.600 | 1.950 | 0.038 |
| **PW set 2** | 1.200 | 1.800 | 2.000 | 2.200 | 2.600 | 2.003 | 0.040 |
| **PW set 3** | 1.400 | 1.800 | 2.000 | 2.200 | 2.400 | 1.973 | 0.040 |
| **MEG: Logarithmic word frequency** |  |  |  |  |  |  |  |
| **Words** | 0.000 | 1.512 | 2.229 | 2.858 | 4.032 | 2.137 | 0.115 |
| **MEG: OLD20** |  |  |  |  |  |  |  |
| **Words** | 1.600 | 1.750 | 1.850 | 1.900 | 2.050 | 1.825 | 0.013 |
| **novel PW** | 1.250 | 1.637 | 1.750 | 1.863 | 2.300 | 1.743 | 0.027 |
| **familiar PW** | 1.250 | 1.600 | 1.725 | 1.863 | 2.100 | 1.717 | 0.026 |
| **MEG: Trigram frequency** |  |  |  |  |  |  |  |
| **Words** | 1.773 | 2.573 | 2.785 | 3.070 | 3.778 | 2.804 | 0.049 |
| **novel PW** | 1.176 | 2.402 | 2.684 | 2.915 | 3.597 | 2.649 | 0.060 |
| **familiar PW** | 1.342 | 2.585 | 2.684 | 2.919 | 3.425 | 2.670 | 0.407 |
| **MEG: Number of syllables** |  |  |  |  |  |  |  |
| **Words** | 1.00 | 2.00 | 2.00 | 2.00 | 2.00 | 1.817 | 0.050 |
| **novel PW** | 1.00 | 2.00 | 2.00 | 2.00 | 2.00 | 1.933 | 0.032 |
| **familiar PW** | 1.00 | 2.00 | 2.00 | 2.00 | 2.00 | 1.95 | 0.028 |
| **MEG: Perimetric complexity** |  |  |  |  |  |  |  |
| **Words** | 5.000 | 5.800 | 6.400 | 6.800 | 7.600 | 6.260 | 0.086 |
| **novel PW** | 5.000 | 5.800 | 6.400 | 6.800 | 8.400 | 6.323 | 0.091 |
| **familiar PW** | 5.200 | 5.800 | 6.200 | 6.800 | 7.800 | 6.327 | 0.627 |
| **MEG: Number of simple features** |  |  |  |  |  |  |  |
| **Words** | 1.400 | 2.000 | 2.200 | 2.400 | 2.800 | 2.160 | 0.038 |
| **novel PW** | 1.200 | 1.800 | 2.000 | 2.200 | 2.400 | 1.983 | 0.040 |
| **familiar PW** | 1.200 | 1.800 | 2.000 | 2.200 | 2.400 | 1.913 | 0.298 |

See Figure S1 for parameter correlations.

#### 2.1.3 Analyses

##### Statistical Modeling

Linear mixed models (LMMs) were used to investigate the three-way and subordinate two-way interactions of each of the four visual and pre-lexical word parameters (see below) with the factors context (i.e., the priming effect of primed vs. unprimed stimuli) and stimulus lexicality (words vs. novel pseudowords) in log-transformed response times, using the lmerTest package (Kuznetsova et al., 2017) of the statistical software package R, version 3.5.3, 2019-03-11 (R Development Core Team, 2008). The model structure is shown in the left panel of Figure 2. In the case of the word frequency parameter, only the interaction with context was included, as an interaction with lexicality is not possible (all pseudowords were assigned a word frequency of zero). Note that in the behavioral study, the factor context was operationalized as the contrast between repeated targets in priming blocks vs. single items in non-priming blocks. Therefore, ‘context’ here represents the effect of the presence vs. absence of contextual information on processing of the target stimulus, whereby only valid contextual information was considered while trials in which the target was preceded by a non-identical prime were discarded (analogous to the MEG experiment; see below). As a consequence, the priming condition had fewer trials (.75 x 60 = 45; minus errors) than the non-priming condition. However, LMMs with crossed random effects are optimal for the analysis of imbalanced data (Baayen et al., 2008). The two-way interaction terms between word parameters (e.g., word frequency) and context were used to determine whether the effect of the respective word parameter was modulated by a predictive context. The three-way interaction with lexicality additionally revealed whether the ‘context by word parameter’ modulations differed between words and pseudowords. This allowed us to assess whether context-based facilitation at the respective level relies on prior knowledge, which is available for words but not pseudowords. Note that we included these interaction terms, i.e., with context and lexicality, for all word parameters within one single LMM. Trials in which errors occurred (9.8 %) were excluded from analyses. We know from previous experience that trial order can have a strong effect on response times and neuronal activation. To explicitly account for this, trial order was included into the LMMs as fixed effect. Participant and item were included as random effects on the intercept. For visualization of partial effects, i.e., effects of a parameter of interest after partialing out all other effects in the LMM, we used the *remef* package (Hohenstein and Kliegl, 2015).

**Figure 2.**
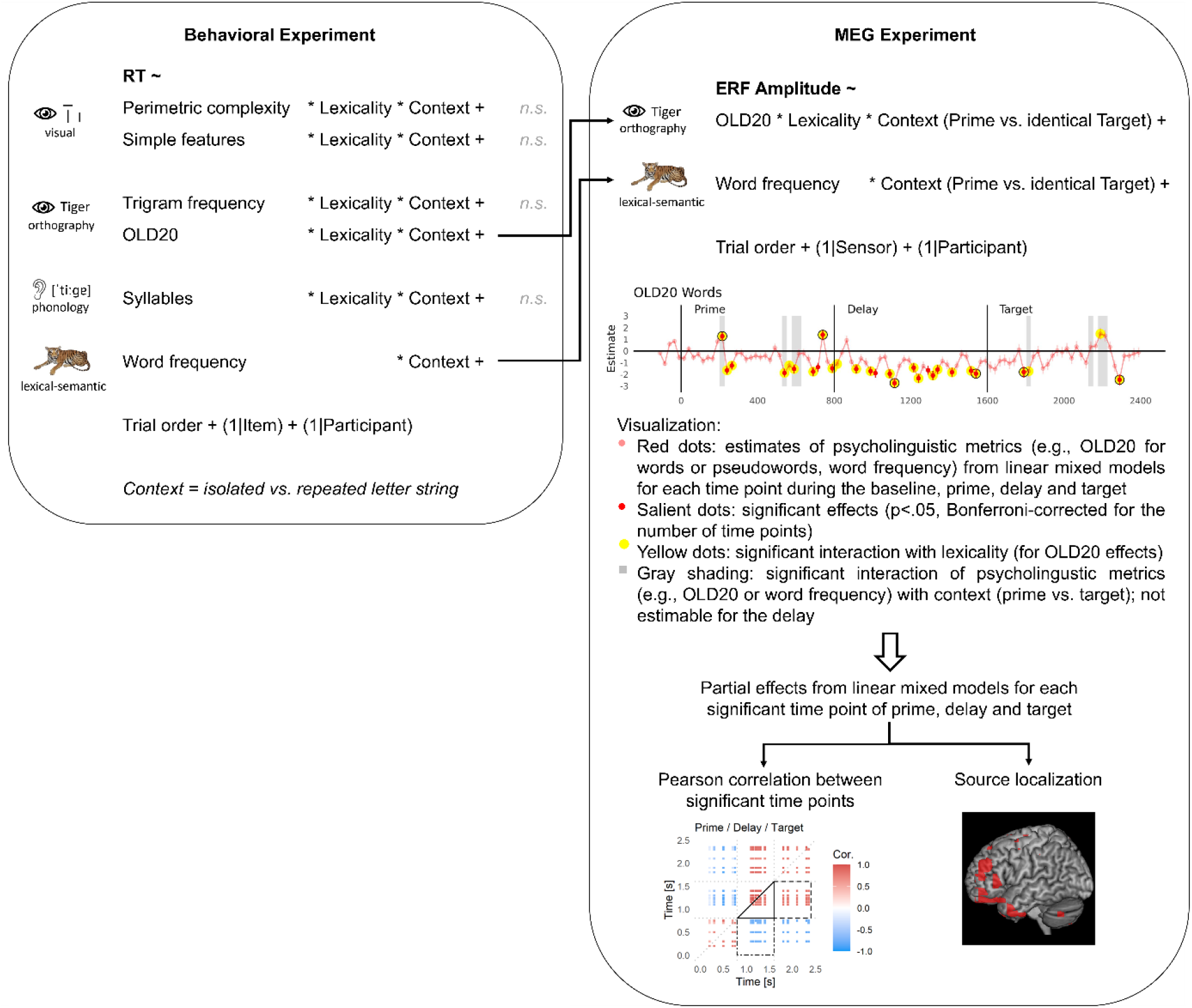
Analys is pipeline for behavioral (top) and MEG data (right). In the behavioral experiment, a linear mixed model assessed the interaction of context (isolated vs. repeated letter strings) with various psycholinguistic metrics associated to visual. orthographic, phonological or lexical-semantic representational levels. In the MEG experiment, linear mixed models were estimated for each time point and included only those psycholinguistic metrics that were found significant in behavior. For each significant effect in the MEG data, partial effects were estimated from the linear mixed models. Pairwise correlations between partial effects were computed across time point. Brain regions underlying the partial effects were identified via source localization. RT = response time, *n.s.* = not significant, ERF = event-related field. • denotes interactions and (1| …) denotes random effects on the intercept.

##### Investigated word parameters

Different descriptors of words were chosen in order to isolate different ‘levels’ of processing a word. *‘Early visual processing’* of a written word depends on the complexity of its physical appearance, which we here characterize, following Chang et al. (2018), using perimetric complexity (describing the density of black pixels in relation to white background which has previously been associated with letter identification efficiency; see Pelli et al., 2006) and the number of simple features that make up a word (i.e., the number of strokes per letter). *‘Pre-lexical processing’* of written words comprises phonological and orthographic processing (e.g., see Carreiras et al., 2014). Phonological processing can be captured by the number of syllables of a word (as syllables reflect sublexical units for sequential phonological processing; e.g., Álvarez et al., 2010; Chetail, 2014), and orthographic processing is captured by the orthographic Levenshtein distance 20 (OLD20, Yarkoni et al., 2008a) and by trigram frequency (e.g., Colegate and Eriksen, 1972; Chen et al., 2015). Lastly, lexical processes of word identification are often associated with word frequency (e.g., Forster and Chambers, 1973; Fiebach et al., 2002), so that logarithmic word frequency (Brysbaert et al., 2011) is included as a parameter representing *‘lexical-semantic processing’*. Pseudowords were assigned a word frequency of zero as they did not appear in the SUBTLEX database. Finally, we included a binary lexicality contrast (words vs. pseudowords) comparing items with and without semantic associations investigating lexical-semantic processing.

##### Model comparisons

In cases with non-significant interaction effects on one or more processing levels, we repeated the analysis with a simpler model in which only the significant interactions and main effects were included. This sparser model was compared to the full model based on the difference in the Akaike information criterion (AIC). The AIC allows comparing models of different complexity (i.e., with more or fewer parameters included). A significantly lower AIC for a more complex model indicates an increase in model fit with the newly added parameter. If the AIC difference is positive or equal, the sparser model has a better fit, and the addition of the new parameter is not advised. Our results and interpretations will be based on the model that includes the set of parameters that lead to an optimal fit.

##### Control analyses

Besides our main analysis of interest described above, we performed two control analyses. First, we re-estimated the LMM with optimal fit while including the familiar as opposed to the novel pseudowords. This allowed us to assess whether the observed lexicality effects are driven by lexical-semantic information, which is available for words but neither pseudoword group. If this is the case, lexicality effects should be observed both for words vs. novel pseudowords as well as for words vs. familiar pseudowords. In contrast, if lexicality effects are based on the different general familiarity with words vs. novel pseudowords, the lexicality effect should be diminished when contrasting words with familiar pseudowords.

The analyses so far were focused on trials of repeated targets and isolated items. Trials of non-repeated targets, i.e., in which prime and target were not identical, were seldom (12.5 % of trials). In a second control analysis, we compared response times between repeated targets, non-repeated targets, and isolated letter strings irrespective of letter string characteristics. This analysis served as a manipulation check to confirm that responses to repeated targets (predictable letter strings) are faster than responses to both non-repeated targets (mispredicted letter strings) and isolated (unpredictable) letter strings.

### 2.2 Magnetoencephalography (MEG) experiment

#### 2.2.1 MEG data

We used data from a previously published repetition priming study (Eisenhauer et al., 2019; data publicly available from https://osf.io/fc69p/), comprising MEG recordings from 38 healthy right-handed native speakers of German (26 females) between 18 and 31 years of age and with normal reading abilities. We here provide a short description of the data acquisition and experimental procedures, and refer the reader to our earlier paper for a detailed description. Data were recorded using a MEG system with 269 operating third-order axial gradiometers (Omega 2005; VSM MedTech Ltd., Coquitlan, BC, Canada) at a sampling rate of 1,200 Hz, applying online filtering between 0.1 and 300 Hz. Procedures were approved by the local ethics committees (University Clinic of Goethe University Frankfurt, application N° 107/15; and Department of Psychology, Goethe University Frankfurt, application N° 2015-229).

The experimental paradigm was similar to the behavioral experiment described above: During MEG acquisition, participants silently read words or pseudowords of five letters length (black on white background; 14 pt at 51 cm viewing distance), which were presented in a repetition priming paradigm with 75 % identical prime-target pairings (repetition trials) and 25 % non-identical prime-target pairings (non-repetition trials; Fig. 1b). Prime and target stimuli were presented for 800 ms each, separated by a delay period of 800 ms during which a string of five hash marks was presented. Primes could either be real words, novel pseudowords (as defined above), or familiar pseudowords (without learned meaning). The pseudoword familiarization procedure and outcomes are described in Supporting Results 2. Each of these conditions consisted of 120 items, i.e., 60 stimuli presented twice. To ensure the participants’ attention to the presented letter strings, 80 additional catch trials were randomly interspersed into the trial sequence, during which the word *Taste* (Engl. *button)* indicated the requirement to press a button. In total, 440 trials were presented across three blocks with a total duration of ∼40 min. 120 trials per letter string condition were considered for analyses of prime and delay intervals whereas 90 items per condition were used for analyses of the target interval (as only 75 % of trials were repetition trials; see also below). Words and both pseudoword groups were matched for OLD20 and number of syllables (see Table 1, also including further parameters, and Fig. S1 in Supporting Information for parameter correlations).

For source localization, structural magnetic resonance (MR) images obtained with a 1.5 T Siemens magnetom Allegra scanner (Siemens Medical Systems, Erlangen, Germany) using a standard T1 sequence (3D MPRAGE, 176 slices, 1 x 1 x 1 mm voxel size, 2.25 s TR, 2.6 ms TE, 9° flip angle) were available for 34 participants (Eisenhauer et al., 2019). The MR images contained fiducial markers for the two auricular MEG head localization coils to improve co-registration with the MEG data.

#### 2.2.2 Preprocessing

Initial preprocessing steps were performed with the FieldTrip toolbox, version 2013 01-06 (http://fieldtrip.fcdonders.nl; Oostenveld et al., 2011) under MATLAB (version 2012b, The MathWorks Inc., Natick, MA). Parallel computations were performed using GNU Parallel (Tange, 2011). MEG data were segmented into epochs of 2,600 ms length, lasting from −200 ms to 2,400 ms with respect to the onset of the prime. Trials contaminated with sensor jump or muscle artifacts, as well as trials in which the head position deviated more than 5 mm from the participant’s average head position across all trials, were rejected, and trials contaminated with eye blink, eye movement, or heartbeat artifacts were cleaned using Independent Component Analysis (ICA, Makeig et al., 1996). See Eisenhauer et al. (2019) and our analysis scripts (https://osf.io/fc69p/) for detailed procedures. After preprocessing, an average of 69 trials (range: 26 to 79) per condition remained for analyses involving both repetition and non-repetition trials (see below), and an average of 51 trials (range: 37 to 106) remained for analyses including only repetition trials. Data were low-pass filtered at 20 Hz and baseline corrected using the time window from −110 to −10 ms; all samples between −110 and 2,400 ms were used for subsequent analyses. To reduce computational costs, the MEG data were down-sampled to 40 Hz.

#### 2.2.3 Linear mixed model analyses

The analysis pipeline for the MEG data is outlined in the right panel of Figure 2. In the light of the variability of previous findings with respect to whether predictive processing during word recognition includes multiple representational levels, we used LMMs for the MEG data analyses. LMMs provide a higher statistical power via single-trial analysis (e.g., Frömer et al., 2018) and inclusion of crossed random effects (Baayen et al., 2008) as opposed to analyses performed on aggregated data. Also, LMMs are being used increasingly for the analysis of electrophysiological data (e.g., Fruchter et al., 2015; Payne et al., 2015).

LMMs were computed on event-related field (ERF) values separately for each time point of the baseline, prime, delay, and target time windows (resulting in one LMM per sampled time point). LMMs were computed in sensor space to reduce complexity (cf. 269 sensors vs. 1,619 source voxels), while a subsequent source localization of significant sensor-level effects (see next section) revealed the underlying brain regions (cf. Manahova et al., 2018; Dijkstra et al., 2018 for a similar procedure). For the LMM analysis of the target time window, we excluded non-repetition trials (in which targets were not predictable) in order to focus on predictable targets only. For the remaining analyses (i.e., concerning prime and delay interval) we included repetition and non-repetition trials as these did not differ between repetition and non-repetition trials. The models were computed using the lmerTest package (Kuznetsova et al., 2017) in R, which allows computing *p-values* for LMMs (Luke, 2017). Resulting *p-values* were Bonferroni-corrected for multiple comparisons to a critical *p-value* of .05/101 = .000495. Following Krakauer et al., (2017), we focused on those effects for which a behavioral correlate was identified in the independent behavioral experiment, i.e., the *pre-lexical/orthographic level of representation* (reflected in a context x OLD20 interaction) and the *lexical-semantic level of representation* (reflected in context x lexicality and context x word frequency interactions; see Results below). Thus, we included as fixed effects only word frequency and the interaction of OLD20 with lexicality (words vs. novel pseudowords). As in the analysis of the behavioral experiment, trial order was included as additional fixed effect of no interest. All participants and MEG sensors were included simultaneously in each model, and random intercepts were estimated for these factors. Based on the logic of parsimonious mixed modeling (Bates et al., 2018), the random effect of item was not included, because including it resulted in a high number of un-estimable models (i.e., non-convergence, unidentifiability, or singluar fit). Still, 13.9 % of models (i.e., of the 101 models calculated for the different time points of the time windows of interest) were un-estimable and thus treated as if they were not significant. Word frequency, OLD20, and trial order were *z*-transformed to enhance the likelihood of convergence and the accuracy of parameter estimates (e.g., Harrison et al., 2018). Post hoc LMMs were computed separately for words and novel pseudowords to resolve interactions with lexicality as well as to obtain the frequency effect for words only; here, 7.9 and 5.0 % of models were un-estimable, respectively.

To explicitly investigate the influence of context (i.e., prime vs. target) on the effects of interest, we combined data from prime and target time windows. We performed this analysis for each of the 32 time points of prime/target presentation, Bonferroni-corrected to a critical *p-value* of .05/32 = 0.0015625. For the interaction of context and lexicality, we computed the full model controlling for effects of the frequency by context interaction and the three-way interaction of lexicality with OLD20 and context (18.8 % un-estimable). For the interaction of context and word frequency or OLD20, we computed separate post hoc models for words and pseudowords (21.9 and 9.4 % un-estimable, respectively). All models included trial order as an additional fixed effect, and participant and sensor as random effects on the intercept.

As for the behavioral data, we performed a control analysis by re-estimating the LMMs while including familiar instead of novel pseudowords. The percentage of un-estimable models amounted to 15.6 % for the analysis including the interaction with context (just prime and target time windows) and 9.9 % for the analysis including all time windows (but no interaction with context). A control analysis including non-repeated trials irrespective of letter string characteristics was already performed in our previous study of the same MEG dataset (Eisenhauer et al., 2019; Fig. 4).

To assess if the strength of pre-activation (i.e., the LMM-based partial effect of psycholinguistic metrics detected as significant in the delay period) is related to the processing strength (i.e., partial effect) of unpredictable as well as predictable letter strings (i.e., prime and target, respectively) at time windows that show a context effect (i.e., a change in neural activation from prime to target), we computed pairwise Pearson correlations between significant time points from the delay interval with prime and target time points showing a significant interaction with context. In the initial analysis, we estimated the effect of interest (lexicality, word frequency, and OLD20 of words and pseudowords) for the specific time points at which the interaction with context was significant. We then aggregated the LMM-based partial effects across participants, sensors, and repeated item presentations before we estimated the correlations. However, in this initial analysis, we noted that almost all correlations were positive, even those between positive and negative effects, indicating that the correlations might be confounded by residual noise not representing the effect of interest. We therefore first performed an LMM analysis modeling the partial effect of the parameter of interest dependent on the parameter as well as on the random effect of item. Remember that we were not able to include the random effect of item in the original LMM analysis as this resulted in a high proportion of un-estimable models. Next, we re-estimated the effect of interest, partialing out the random effect of item. We then again performed the correlation analysis as described above, using the re-estimated effects. Indeed, in this analysis the sign of the correlation effects represented the sign of the observed effects, i.e., positive effects correlated negatively with negative effects. Obtained *p-values* were Bonferroni-corrected for the number of computed correlations (which could differ between the four psycholinguistic metrics). E.g., for lexicality, six time points were found significant during the delay, and seven time points showed a significant interaction with context, resulting in 190 correlations and consequently a Bonferroni-corrected critical *p-value* of .05/190 = 0.00026.

#### 2.2.4 MEG source localization

We estimated source locations from the ERF difference of words vs. novel pseudowords, high vs. low-frequency words, and high vs. low OLD20 (the latter separately for words and pseudowords). Word frequency and OLD20 contrasts were based on median splits to increase the signal-to-noise ratio compared to single-trial analysis (e.g., Gross et al., 2013). As time windows for source analysis we selected the significant time points of lexicality effects, word frequency effects (words only), and the OLD20 effects for words and pseudowords from the LMM analysis (see previous sections). Source localizations of effects obtained in the LMM analysis were conducted on the effects of interest after correcting for all other sources of variance in the model. To this end, we simulated the respective effect of interest from the LMM for each sensor in a way that controls all other effects, using the *remef* package (Hohenstein and Kliegl, 2015). This source localization procedure is optimal when for each time point, all data from all sensors is included in one model, which is possible since we treated the sensors as a random effect. The simulated data from the LMM projects the effect of interest onto the sensors after correcting for all confounding variables, and these simulated sensor data are then subjected to standard source reconstruction procedures as described below. This ‘preprocessing’ procedure for source localization is a central feature of our LMM analysis and currently, to our knowledge, not used elsewhere.

Source localization was performed using linearly constrained minimum variance (LCMV) spatial filters (Van Veen et al., 1997) in FieldTrip, Version 2016 10-24, for those 34 participants for whom anatomical MR images were available. The procedure closely followed our earlier study (Eisenhauer et al., 2019) and is based on Manahova et al. (2018; see also Dijkstra et al., 2018; Mostert et al., 2018). Anatomical MR images were warped to an MNI-space template comprising 1,619 voxels within the brain (10 mm resolution). Two-dimensional dipole moments were computed for each voxel location. A single shell forward model of the inner surface of the skull (Nolte, 2003) was used for lead field computation. The data covariance was regularized with a shrinkage parameter of 0.01. The localization approach comprised a permutation procedure with 1,000 analyses on shuffled data to account for noise. Please refer to Eisenhauer et al. (2019) for a detailed description of the permutation procedure. Finally, the signal of each source location was normalized by its variance to counter the depth bias.

For visualization, LMM-based source localizations thresholded at 50 % of the peak activation across all time windows included in the respective contrast, as well as thresholded at 90 % of the individual peak of the respective time point, were plotted on cortical surfaces using MRIcron (Rorden and Brett, 2000). In order to make the OLD20 source activation strengths comparable between words and pseudowords, both were thresholded at 50 % of the peak activation for words. Also, we visualized source overlaps between the identified sources of prime, delay, and target, based on both the 50 and the 90 % threshold. Brain regions were identified from the MNI coordinates of the ≤3 most robust sources per contrast using the AtlasReader (Notter et al., 2019) in Python.

### 2.3 Code and data accessibility

The analysis code used for behavioral and MEG data analyses, as well as the behavioral data, are publicly available from https://osf.io/c7s2k/. The MEG and MRI data are publicly available from https://osf.io/fc69p/.

## 3 Results

### 3.1 Behavioral experiment: Identifying context-based facilitation effects

In the data from the behavioral experiment, we investigated the influence of a predictive context (i.e., prime absent vs. present) on visual, orthographic, phonological, and lexical-semantic processes during visual word recognition. These different ‘levels’ of processing were approximated in a linear mixed model (LMM) analysis of response times by various psycholinguistic metrics. We thus modeled the influence of context on effects of (i) perimetric complexity and number of simple features of the stimuli (visual processing), (ii) OLD20 and trigram frequency (orthographic processing), (iii) number of syllables (phonological processing), and (iv) word frequency as well as lexicality (words vs. novel pseudowords; lexical-semantic processing).

As our hypotheses were focused on the interactions of these stimulus characteristics with context-dependent facilitation, we will in the following primarily report results involving context effects; full results of the LMMs can be found in associated Tables. In general, processing of expected items (i.e., targets preceded by an identical prime stimulus) was facilitated, i.e., they were responded to faster than unprimed items (significant main effect of context: p < 2e-16; Table 2). This context-dependent facilitation interacted with parametric predictors representing orthographic (i.e., OLD20) and lexical-semantic (i.e., word frequency and lexicality) but not visual or phonological processing (Table 2; see also Table S1 for the results of the full model). In order to implement an adequate statistical model, we removed the non-significant visual, orthographic (i.e., trigram-frequency), and phonological predictors from the full model and re-estimated the model. This reduced model is presented in Table 2; an explicit model comparison indicated that the Akaike Information Criterion (AIC) of this simpler model was 21 points lower than that of the full model (*chi-square* = 11.051, *df* = 16, *p* = 0.806). Given that these two models did not differ statistically, the simpler model should be preferred (see Methods section).

**Table 2.** Results of the linear mixed model (LMM) investigating context effects (repeated target vs. isolated presentation) on word frequency, OLD20, and lexicality in behavioral response times

|  | *FE* | *SE* | *t* | *p* |
| --- | --- | --- | --- | --- |
| Context | **-0.278** | **0.013** | **-21.951** | **< 2e-16** |
| Word frequency | **-0.047** | **0.014** | **-3.312** | **0.001112** |
| Lexicality | -0.019 | 0.027 | -0.715 | 0.476 |
| OLD20 | **-0.013** | **0.006** | **-2.078** | **0.038** |
| Trial order | **-0.025** | **0.002** | **-10.03** | **< 2e-16** |
| Word frequency x Context | **0.039** | **0.012** | **3.316** | **0.000918** |
| Lexicality x Context | **-0.131** | **0.023** | **-5.601** | **2.19e-08** |
| OLD20 x Context | **-0.018** | **0.007** | **-2.596** | **0.009450** |
| OLD20 x Lexicality | 0.007 | 0.011 | 0.603 | 0.547129 |
| OLD20 x Context x Lexicality | **0.035** | **0.011** | **3.233** | **0.001230** |

With respect to *orthographic processing*, a significant 3-way interaction between context, lexicality (words vs. novel pseudowords), and OLD20 (p = .001; Table 2; see also Tables S2 and S3 for resolving this interaction post hoc via two-way interactions) could be resolved to show significant negative effects of OLD20 on response times for novel pseudowords, with faster decision times particularly for expected pseudowords that have a high orthographic similarity to words (i.e., low OLD20; see Fig. 3a, Table 3, and Table S2). For words, we observed descriptively a negative effect of OLD20 for words presented in isolation and a positive effect for primed words (Fig. 3a), which resulted in a significant context by OLD20 interaction for words (see post hoc statistics in Table S2). This interaction could, however, not be further resolved statistically in post hoc analyses (Table 3).

**Figure 3.**
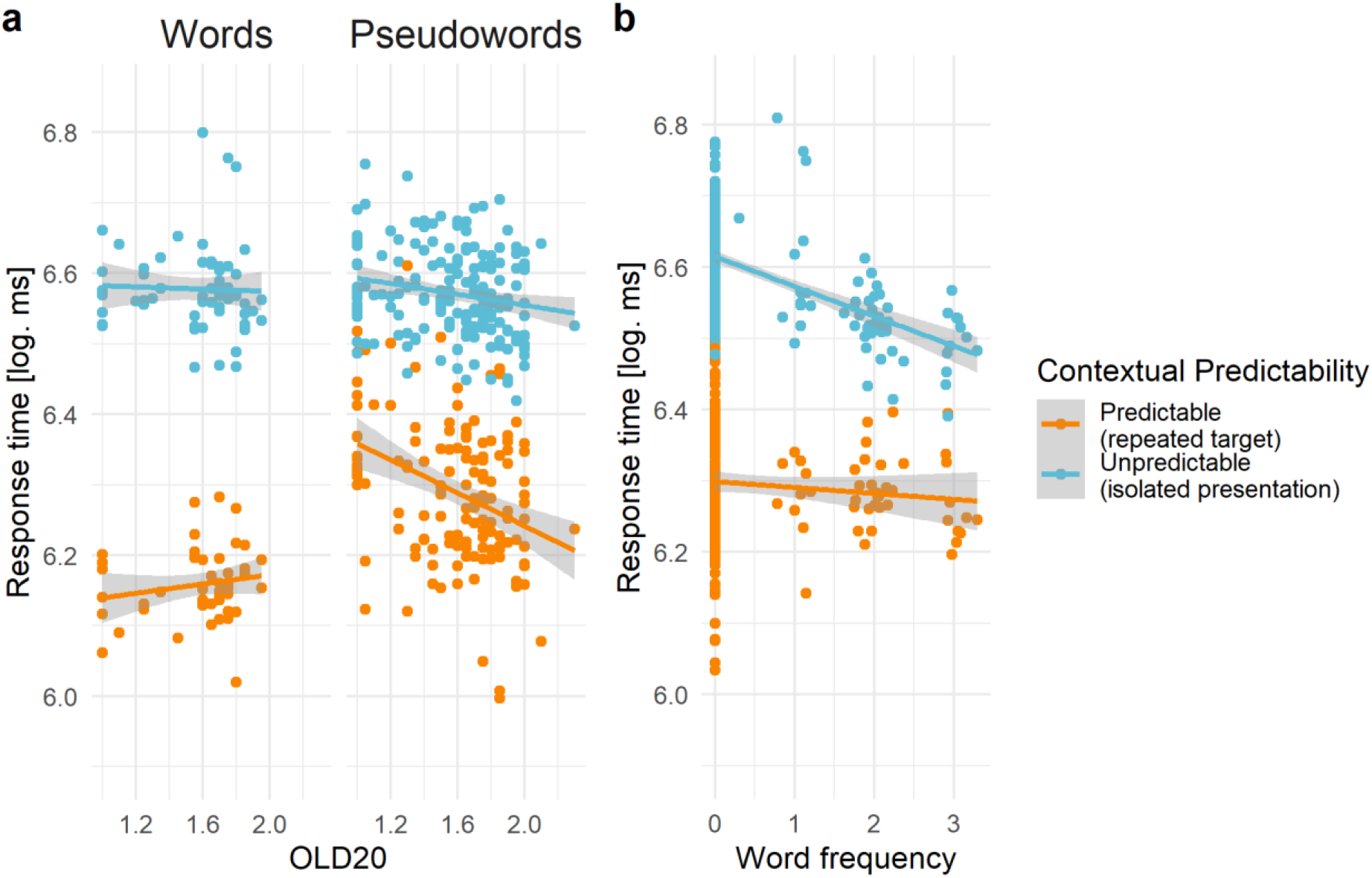
Behavioral results reflecting context-dependent facilitation. Comparison of response times representing (a) orthographic (OLD20 effect separated for words and pseudowords) and (b) lexical processing (word frequency effect) when stimuli were either predictable from context (i.e., repeated targets; orange lines and dots) or not (isolated presentation; blue lines and dots). Note that lower OLD20 values reflect higher word likeness, and, thus, orthographic familiarity. Displayed are logarithmic response times that represent the partial effects estimated from linear mixed models. Dots represent items averaged across participants.

**Table 3.**
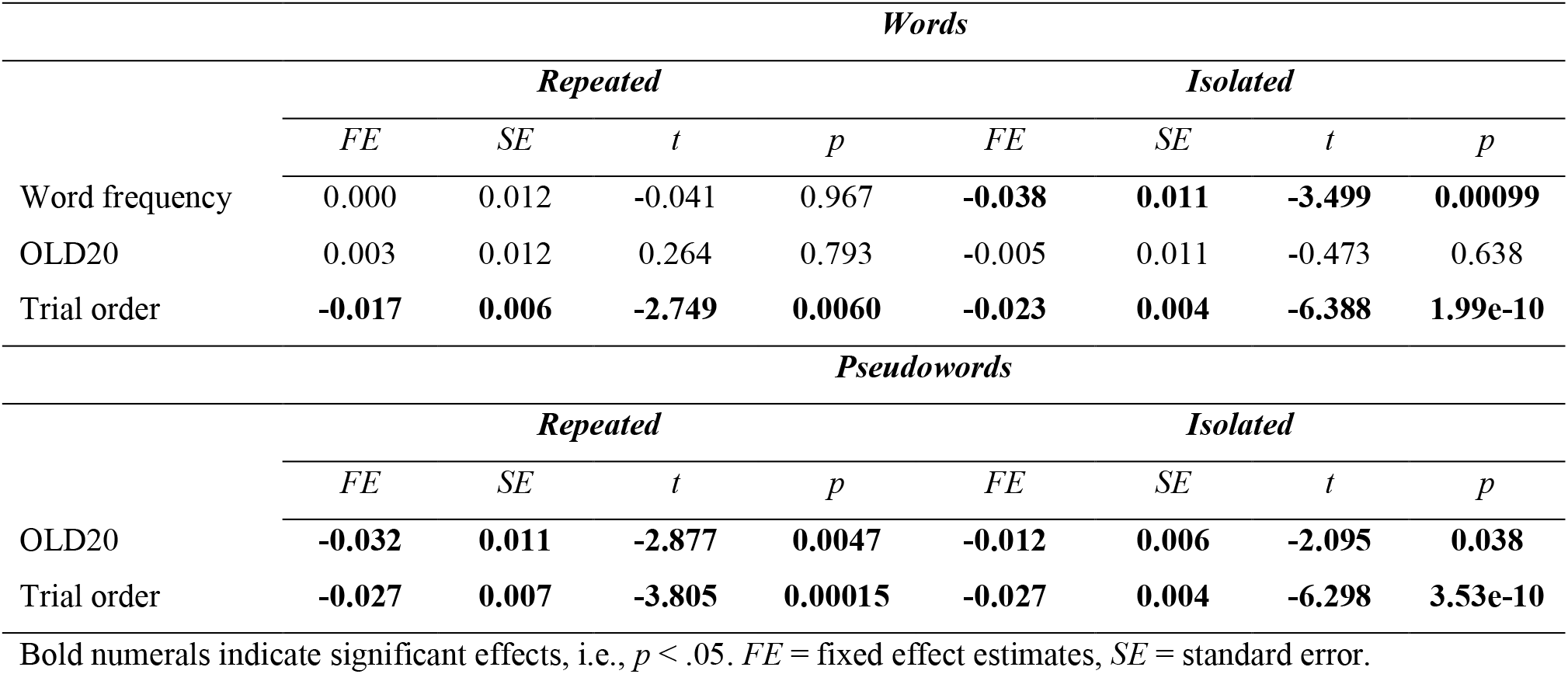
Results of post hoc linear mixed models (LMMs) investigating word frequency and OLD20 effects on behavioral response times separately for repeated and isolated words and pseudowords

With respect to *lexical-semantic processing*, a significant interaction between context and word frequency (p < .001; Table 2) provided evidence for context-dependent facilitation of lexical-semantic processing. Post hoc evaluation showed significantly faster decision times to more frequent relative to less frequent words when presented in isolation (i.e., when not predictable), but no significant frequency effect for primed words (given generally faster responses; Fig. 3b; see also Table 3 for word frequency effects of words only, and Table S3 for word frequency effects combined across words and novel pseudowords, for which a word frequency of zero was assumed). Lexical-semantic processing as one locus of predictive effects is also supported by the observation of a context by lexicality interaction (p < .001; Table 2), which shows that the effect of context-dependent facilitation is generally much stronger for words than for novel pseudowords (see also Fig. 3a; compare also Eisenhauer et al., 2019). In addition, this was confirmed in a control analysis revealing stronger context-based facilitation effects for words than for familiar pseudowords (see Supporting Results 3). Finally, our manipulation check confirmed faster response times not only for repeated targets vs. primes, but also for repeated targets vs. non-repeated targets irrespective of psycholinguistic variables (see Supporting Results 4). To summarize, these behavioral results suggest that context-based facilitation involves processes at the pre-lexical orthographic and the lexical-semantic stages of visual word recognition, while no evidence was found for context effects at the level of visual or pre-lexical phonological processing. As a consequence, we explore the neurophysiological data with respect to neural signatures of predictive context effects and pre-activation effects of OLD20 (orthographic similarity), word frequency, and lexicality.

### 3.2 Investigation of orthographic and lexical-semantic representations in brain data

MEG-measured event-related fields elicited by words and pseudowords of the present dataset were investigated in depth in our earlier publication. The previous findings revealed a context effect of lower neuronal activation for repeated targets (i.e., predictable letter strings) in comparison to both non-repeated targets (i.e., mispredicted letter strings) and primes (i.e., unpredictable letter strings) in left-lateralized language network regions (Eisenhauer et al., 2019; Fig. 4; https://www.eneuro.org/content/eneuro/6/2/ENEURO.0321-18.2019/F4.large.jpg). This finding confirms that our context manipulation was successful in facilitating the processing of predictable letter strings by reducing the required neuronal resources. In addition, the context effect of repeated targets vs. primes was stronger for words compared to both novel and familiar pseudowords within the left anterior temporal cortex (Eisenhauer et al., 2019; Fig. 5H-J https://www.eneuro.org/content/eneuro/6/2/ENEURO.0321-18.2019/F5.large.jpg), indicating context-based facilitation in particular at the lexical-semantic level.

**Figure 4.**
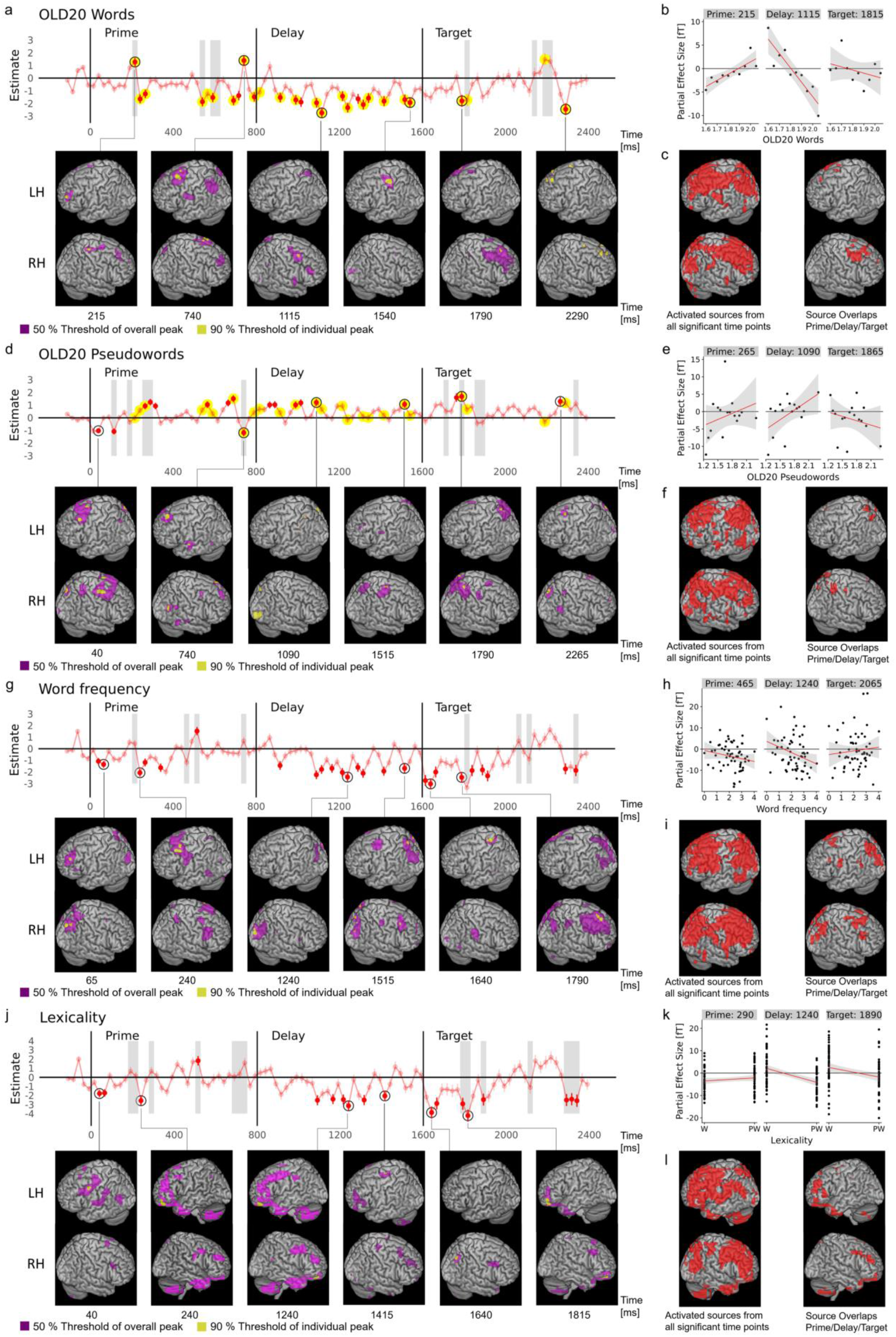
Effects of orthographic and lexical-semantic information across time and their brain localization. (a) Effects of OLD20 during word processing. Displayed are the estimates of linear mixed models ± SE across time, including baseline, prime, delay, and target time windows. The salient red dots represent significant time points, excluding un-estimable models. Gray shading represents time points of a significant interaction between OLD20 and context (prime vs. target). Yellow circles represent time points of a significant interaction between OLD20 and lexicality (words vs. pseudowords; compare panel (b)). Black circles mark time points for which source localizations are shown. Source topographies display sources thresholded at 50 % of the overall peak (maximal activation across all time points; violet), as well as thresholded at 90 % of the individual peak per time point (yellow) within the left (LH) and right (RH) hemisphere. (b) Scatterplots depict model-estimated partial effect sizes for three exemplary time points from prime, delay, and target time windows. In case of prime and target, a time point showing a significant interaction between the respective parameter and the prime/target difference was selected. (c) Activated sources from all significant time points (left) and overlapping sources during prime, delay, and target time windows (right), based on both the 50 % threshold of the overall peak as well as the 90 % threshold of the individual peak. (d,e,f), (g,h,i), and (j,k,l) show corresponding results for OLD20 during pseudoword processing, word frequency (for words only), and lexicality, respectively. To facilitate comparability of OLD20 source activation strengths for words and pseudowords, the 50 % threshold of OLD20 source activations for words was used for pseudowords as well. See Table S10 for source coordinates.

**Figure 5.**
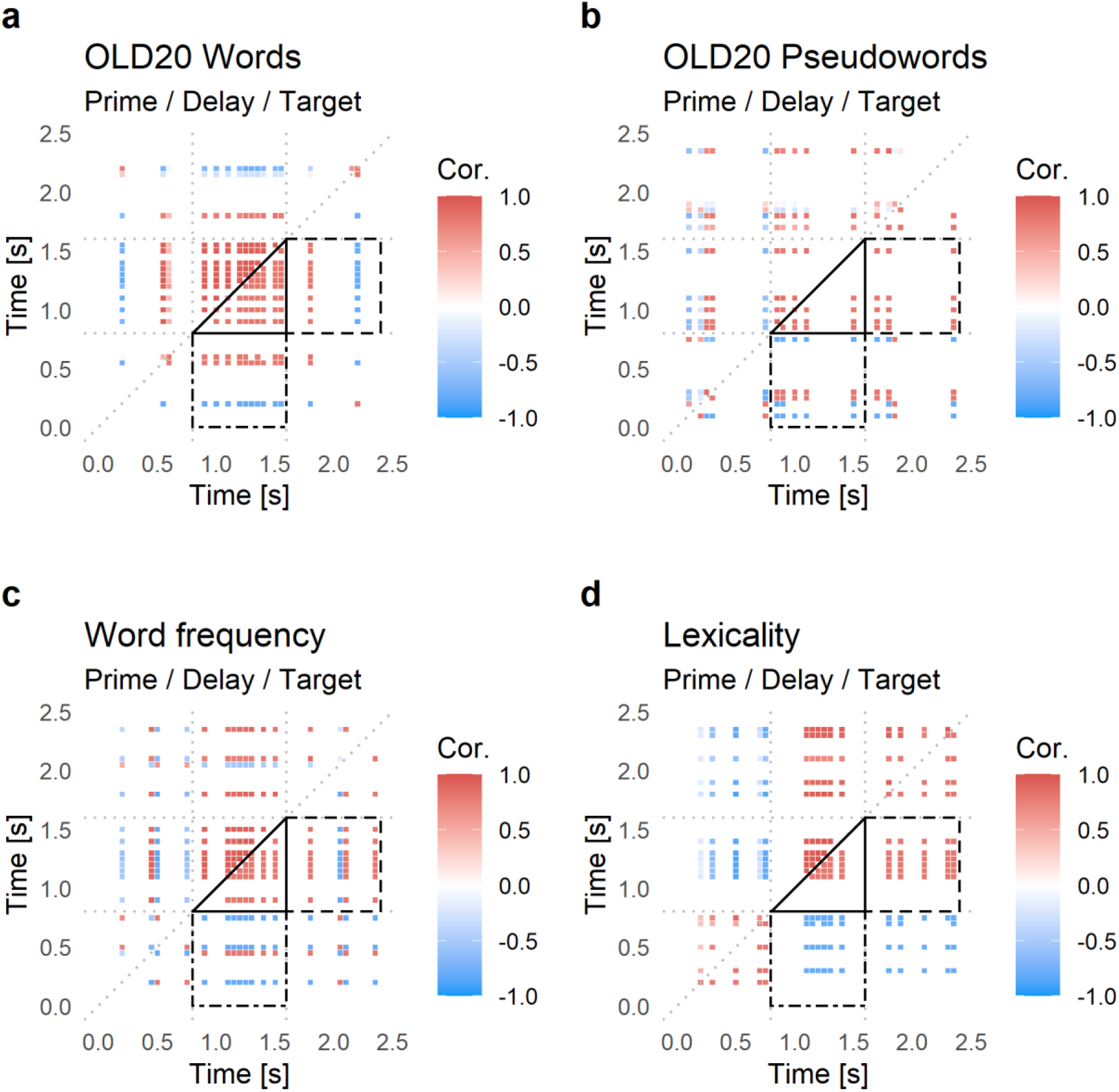
Correlations between LMM-based partial effects across prime, delay, and target time points for (a) OLD20 effects for words, (b) OLD20 effects for pseudowords, (c) word frequency, and (d) lexicality. For the delay, correlations were computed for all time points showing a significant effect of the respective letter string characteristic, while for prime and target, time points at which effects changed between prime and target (as reflected in a significant interaction of the characteristic with the context effect) were selected. The time-resolved correlations are down–sampled by a factor of 10. Significant correlations are shown below the diagonal, while all correlations are shown above the diagonal. Black triangles mark within-delay correlations, while the dot-dashed black square marks prime-delay correlations and the dashed line black square marks delay-target correlations.

The present work goes beyond this initial report and uses LMMs to directly assess (i) context effects during stimulus processing and their effect on orthographic and lexical-semantic levels of linguistic processing (represented by the psycholinguistic metrics OLD20, word frequency, and lexicality), and (ii) their dependency on pre-activation of orthographic and lexical-semantic representations during the prime-target interval. To this end, we first characterize the temporal dynamics of the effects of OLD20 (reflecting orthographic representations), as well as word frequency and lexicality (both reflecting lexical-semantic representations) by individually analyzing each time point across the entire trial. We then directly compare respective effects identified during processing of prime and target, to quantify context (i.e., repetition) effects at the respective level of representation. Third, we explore pre-activation by testing for the presence of orthographic and lexical-semantic effects in the prime-target delay. In detail, we assess whether these signatures of pre-activation correlate with activation during prime and target processing, primarily to determine how the processing of expected stimuli mechanistically depends on context-dependent pre-activation.

Analogous to the statistical approach used for analyzing the data from the behavioral experiment, we investigated the effects of the three psycholinguistic metrics of interest on brain activation using linear mixed models (LMMs). LMMs model data at the level of individual trials, and thus result in high statistical power (Matuschek et al., 2017). Indeed, while we also investigated brain activation using a multivariate pattern decoding approach (see Supporting Results 5), which has been established in previous work for investigating brain responses in the absence of external stimulation (e.g., Simanova et al., 2015; Heikel et al., 2018), a power analysis indicated higher power and more reliable effect size estimates for LMMs in contrast to multivariate pattern decoding (see Supporting Results 6). Importantly, LMMs can estimate the effects of interest while controlling for confounding variables. Note that LMMs are calculated for every sample time point; in Figure 4, we visualize the LMM estimates, reflecting the modulation of the ERF amplitude at the respective time point. Higher absolute estimate values indicate a steeper regression slope, i.e., a stronger modulation of neuronal activation by the respective parameter. The slopes were thus our effect size measure of the respective stimulus characteristics in neuronal activation. In detail, high effect size values can be interpreted as a strong representation of the respective concept in neuronal activation as they indicate that neuronal activation differentiates between letter strings of higher vs. lower parameter values. In the presence of a negative ERF component, a positive estimate indicates that letter strings with a higher value on the predictor variable (e.g., word frequency) have a less negative-going ERF, and vice versa for negative estimates. The scatterplots in Figure 4 visualize partial effects for exemplary time points, i.e., they represent the data (averaged across participants) and the regression slopes for an effect of interest after partialing out all other effects from the LMMs.

#### Effects of orthographic representations

LMMs revealed a significant interaction between OLD20 and lexicality (words vs. novel pseudowords) – corresponding to the interaction found in our behavioral data – in several time windows (Bonferroni-corrected; marked with yellow in Fig. 4a,d). The interaction was found (i) during presentation of the prime between 215 and 265 ms and between 540 to 790 ms, (ii) during the prime-target delay at multiple time points between 15 and 740 ms after prime offset (corresponding to 815 to 1540 ms relative to prime onset), and (iii) during presentation of the target, i.e., between 190 and 215 ms, as well as 590 and 690 ms after target onset (corresponding to 1790 to 1815 and 2190 to 2290 ms relative to prime onset). The interaction resulted from predominantly negative effects for words (i.e., greater negative-going MEG activity the higher the OLD20 parameter, i.e., the less word-like a word was; see detailed effects in Fig. 4a) and mainly positive effects for novel pseudowords (Fig. 4d). The opposite effects of orthographic similarity (OLD20) on MEG responses elicited during word vs. pseudoword processing are also documented by a negative correlation of the two time courses of the OLD20 effect estimates (r = -.29; t(99) = −3.0; p = .003). Separate investigations of OLD20 effects for words vs. novel pseudowords revealed an early negative OLD20 effect for pseudowords (i.e., at 40 ms) during prime presentation (Fig. 4d). In contrast, the OLD20 effect for words reached significance considerably later, at 215 ms during prime presentation (Fig. 4a). During target processing, OLD20 effects reached significance shortly before 200 ms, with a slightly earlier onset for pseudowords (165 ms) vs. words (190 ms). In the delay period, significant effects were more abundant for words (40.6 % of investigated time points) vs. pseudowords (18.8 %). When comparing the three time windows, words showed overlapping source activations in bilateral frontal and pseudowords in bilateral parietal regions (Fig. 4a,c vs. d,f). When quantifying the contextually-mediated effect of orthographic processing by directly contrasting OLD20 effects during prime and target processing, significant differences emerged at 215, 540, and 590 to 615 ms for words and at 115, 190, 265 to 290, and 740 ms for pseudowords, reflecting a reversal of the effect direction from prime to target at these time points (i.e., from negative to positive or vice versa; marked by gray shading in Fig. 4a and Fig. 4d; see also Fig. 4b and Fig. 4e).

To examine the influence of orthographic pre-activation (i.e., the OLD20 effects) on target processing, we correlated significant effect size estimates from the delay with all timepoints showing significant interactions with context (i.e., that show a relevant change from prime to target). First, we expect that time points within the delay period correlate positively when neuronal pre-activation is maintained stable across time (cf. King and Dehaene, 2014). Positive correlations between prime and delay might either reflect a ‘spill-over’ from prime processing, or a re-activation of representations activated while processing the prime, potentially facilitating the processing of the identical target. We expect a negative correlation between delay and target, when the higher delay activation results in more efficient target processing (i.e., as assumed by predictive coding; Friston, 2005). In contrast, if the observed reversal of the effect direction from prime to target already takes place during the delay, this could indicate that more extensive processing of the prime leads to more efficient pre-activation and target processing.

Within the delay period (see black triangles in Fig. 5a,b), significant effects of OLD20 are highly correlated, but more sustained across the delay for words than for novel pseudowords (Fig. 5a,b). This indicates a stable neuronal activation across the delay period for words, while pseudowords showed less orthographic pre-activation during the delay (see above). From prime to delay, the majority of significant correlations for words was positive (74.0 %), while significant correlations for novel pseudowords did not indicate a stable pattern as only 44.4 % were positive (see the dot-dashed black squares in Fig. 5a,b). Delay-target correlations (see the dashed line black squares in Figs. 5a,b) represent relationships between orthographic pre-activation and processing of the expected item. For words, early target activation (215 ms) was correlated positively, while late target activation (590 to 615 ms) was correlated negatively with orthographic delay activation (Fig. 5a). For novel pseudowords, the target activation was consistently positively correlated with the delay activation (Fig. 5b). To summarize, these findings indicate more sustained pre-activation of orthographic information for predicted words than pseudowords. However, pre-activated information more consistently resembled target effects for pseudowords compared to words; for words, early target effects resembled delay activation, while late target effects indicated stronger facilitation with stronger pre-activation.

#### Lexical-semantic representations: Effects of word frequency

As for OLD20, we found significant effects of word frequency (investigated only for the word condition) at several time points across the prime, delay, and target time windows (Fig. 4g,h). Throughout the entire trial, we found predominantly negative effects (i.e., more negative/less positive activation for higher frequency words), with the exception of one single positive effect at 515 ms after prime onset. Specifically, negative effects were found in the time ranges of 40 to 65 ms and 240 to 340 ms after prime onset. During the prime-target delay, we observed negative effects at 115 ms and at several time points later in the delay, in the time range from 290 to 715 ms (i.e., 1090 to 1515 ms relative to prime onset). During the target presentation, significant negative frequency effects were found in time windows starting earlier when compared to the prime interval, i.e., between 15 and 65 ms. In addition, negative frequency effects were found in the range from 190 to 315 ms (1615 to 1665 ms and 1790 to 1815 ms relative to prime onset), but also at late time points (690 and 740 ms, i.e., 2290 and 2340 ms relative to prime onset). Sources of the word frequency effects across the three time windows were localized to frontal and parietal brain regions (Fig. 4i).

To directly assess context-dependent word frequency effects, we compared word frequency effects between prime and target – statistically assessed as a context by frequency interaction. We identified significant interactions at 215, 465, 515, and 740 ms after stimulus onset (gray shading in Fig. 4g), indicating a reversal of the effect direction from prime to target. As for orthographic representations, we assessed correlations among effect strengths involving all time points with significant differences in frequency effects between prime and target (i.e., context x frequency interactions) and all significant time points from the delay period. Analogous to orthographic effects in the word condition, we found highly correlated effects within the delay period (Fig. 5c), suggestive of a consistent neuronal pre-activation of word frequency-related information. We found predominantly negative correlations when prime and delay periods were compared and predominantly positive correlations between pre-activation effects in the delay and target processing (75.0 % of significant correlations each). These findings suggest a reversal of the word frequency effect direction from prime to delay but relatively consistent neuronal activation patterns throughout the delay and target periods.

#### Lexical-semantic representations: Effects of lexicality

The time course of effect estimates for the second psycholinguistic metric of lexical-semantic processing, i.e., stimulus lexicality (words vs. novel pseudowords), was very similar to that of word frequency (r = .94; t(99) = 26.7; p < .001; compare Figs. 4g vs. j). In detail, during the presentation of the prime, negative lexicality effects (reflecting more negative activation for pseudowords relative to words) reached significance at 40 to 65 and at 240 ms, and we found a positive effect at 515 ms post-stimulus onset. During the delay, significant negative lexicality effects were found for several time points in the range from 290 to 615 ms (corresponding to 1090 to 1415 ms relative to prime onset). During the presentation of the target, significant negative effects were found at 40 to 65, between 190 and 290 ms, and from 690 to 740 ms.

Context by lexicality interaction effects partly overlapped with context interactions involving word frequency: Lexicality effects (words vs. novel pseudowords) differed between prime and target at 190 to 215, 290, 515, and 690 to 740 ms after stimulus onset (gray shading in Fig. 4j; see also Fig. 4k). For the lexicality contrast of words vs. pseudowords, the interaction with the context effect of prime vs. target, as well as the lexicality effects during the delay, reached significance at time points similar to the words vs. novel pseudowords contrast (see Supporting Fig. S3). This finding indicates that the lexicality effects likely capture differences in lexical-semantic representations as opposed to general familiarity differences between the letter string groups (e.g., see Gregorova et al., 2021 for a finding indicating domain generality of lexical access).

In the correlation matrices for the lexicality contrast of words vs. novel pseudowords we found a very clear picture (Fig. 5d). First, all effects within the delay were highly positively inter-correlated. Again, the prime and delay period effects showed negative correlations and delay and target period effects showed positive correlations. When localizing the lexicality effect, we found similar regions across prime, delay, and target time windows, located within frontal, temporal pole, and cerebellar regions. Thus, despite similar time courses, there is an apparent dissociation from the frequency effect (cf. Fig. 4g,i vs. j,l).

## 4 Discussion

We investigated the role of visual, orthographic, phonological, and lexical-semantic representations for context-dependent (i.e., predictive) facilitation of visual word recognition. Consistent with abundant previous literature (e.g., Tulving and Schacter, 1990; Kutas and Federmeier, 2011), context-dependent facilitation is reflected in faster response times and an amplitude reduction in the event-related fields (ERFs) for repeated relative to initial stimulus presentation (see Eisenhauer et al., 2019, Fig. 4, for ERFs from the present dataset). Our behavioral data indicate that orthographic and lexical-semantic, but not visual or phonological processing, interact with context-dependent facilitation. The MEG data show differential effects of orthographic word similarity, word frequency, and lexicality on prime vs. target processing. Importantly, all three parameters also modulated brain activity in the delay interval, providing evidence for pre-activation of orthographic and lexical-semantic representations in predictive contexts. This finding is crucial as the majority of previous studies indicating predictive pre-activation in visual word recognition had not dissociated at what representational level information is pre-activated (e.g., Dikker and Pylkkänen, 2013; Bonhage et al., 2015; Wang et al., 2017). Consequently, our findings further support the previous evidence for predictive pre-activation of lexical-semantic representations (Fruchter et al., 2015; Wang et al., 2020) by showing that lexicality and word frequency are represented in neuronal activation patterns prior to predictable letter strings. In addition, our results extend previous findings by providing the first direct evidence for a pre-activation of orthographic information. A change in the direction of estimated effects reflected a change in the quality of pre-activated lexical-semantic representations from prime to delay, while delay and target effects were positively correlated.

### 4.1 Context-Dependent Facilitation of Visual Word Recognition

Contextual facilitation was modulated by orthographic word similarity. As expected (e.g., Balota and Chumbley, 1984; Fiebach et al., 2007), we found fast non-word responses when orthographic familiarity is low. Predictability amplified this effect and reversed the weak OLD20 effect for words, suggesting an interactive role of orthographic processing and context-dependent facilitation. Regarding lexical-semantic processing, we found stronger behavioral priming effects for words than pseudowords (replicating, e.g., Fiebach et al., 2005) and reduced word frequency effects for expected words (replicating, e.g., Forster and Davis, 1984; Kinoshita, 1995). As word frequency effects are typically interpreted as reflecting differences in the ease of lexical access (e.g., Coltheart et al., 2001), this (together with the generally faster target response times) indicates that the effort of accessing meaning is reduced in highly predictive contexts. Consistent with current literature, we found no evidence for context-based facilitation of visual (e.g., Slattery and Yates, 2018) or phonological processes (e.g., Nieuwland et al., 2018).

### 4.2 Neurophysiological Mechanisms of Context-Dependent Facilitation

Pre-lexical orthographic and lexical-semantic representations modulated brain activation during prime, delay, and target intervals. Orthographic familiarity elicited negative and positive effects for words and pseudowords, respectively, at around 200 and 500 ms. Lexical-semantic effects were identified earlier (before 100 ms) and around 250 ms, both showing a negative modulation (e.g., more positive activation for words than pseudowords) independent of predictability. This result (and its localization to frontal brain areas) replicates early MEG effects obtained with similar stimuli (e.g., Wheat et al., 2010; Woodhead et al., 2014). These early effects might reflect top-down constraints from (higher) lexical-semantic to lower representational levels (e.g., Price and Devlin, 2011; Carreiras et al., 2014). A similar proposal for object recognition (Bar et al., 2006) suggests an initial coarse processing of visual stimuli, which can account for very early top-down effects. According to that model, low spatial frequency components are extracted very early from the visual input and are assumed to be transmitted very rapidly to frontal cortex, triggering higher-level representations that in turn constrain the subsequent more elaborate visual processing of the input (based on high spatial frequency input). The present data, however, provide no insights into how such ultra-fast processing of word semantics might be mechanistically implemented, given that the surface form of words provides no cues for their semantic meaning.

Context-dependent changes of representations (i.e., from prime to target) started at ∼100 ms post-stimulus onset for orthographic (in pseudowords) and ∼200 ms for lexical-semantic representations. These prime-target differences also involved qualitative changes, with larger activations for words vs. pseudowords at the prime and the inverse effect at the target. These findings suggest that context-dependent facilitation is implemented at orthographic and lexical-semantic processing levels (e.g., Almeida and Poeppel, 2013; Brothers et al., 2015).

Orthographic as well as lexical-semantic effects during the delay period emerged several hundred milliseconds before target onset (cf., e.g., Dikker and Pylkkänen, 2013; Gastaldon et al., 2020; for similar time windows of general word pre-activation effects; i.e., without resolving the level at which information was pre-activated). These effects were consistently positively correlated during the delay period, indicating the high stability of activated representations. Effects elicited during the delay reflect, by necessity, either a ‘spill-over’ from processing the prime or predictive pre-activation of target-associated linguistic representations. However, we suggest that the observed delay effects reflect predictive pre-activation for the following reasons: (i) Lexicality and word frequency effects during prime processing were mainly anticorrelated with delay effects, which is incompatible with spill-over as this would cause similar (i.e., positively correlated) neuronal activation patterns during prime and delay. (ii) Spill-over is expected to occur shortly after prime offset, with a sustained representation activated both at the end of prime presentation and the beginning of the delay. Still, no such pattern was observed for lexicality and word frequency. Rather, no significant effects were found during the transition from prime to delay for a period of 575 and 400 ms, respectively. For OLD20 effects of words and pseudowords, correlations between prime and delay were partly positive (74 % for words and 44.4 % for pseudowords), and the time period of non-significant effects at the transition from prime to delay was comparatively short (i.e., 125 ms for both conditions). Nevertheless, the direction of OLD20 effects changes at the end of prime presentation and again at the beginning of the delay, indicating that representations are not sustained over a longer period while transiting from prime to delay. This, however, is not compatible with a spill-over account of the present findings. In contrast to the anticorrelation of lexical-semantic effects during the delay with effects during prime presentation, delay and target effects were positively correlated. This finding indicates qualitative changes from prime to delay and pre-activation of target-associated representations. Early orthographic effects on word primes were anticorrelated with later prime effects (as well as delay and most of the target representations), suggesting that orthographic representations for words are relatively quickly transformed into preparatory codes (i.e., after lexical access which is achieved before 500 ms post-stimulus onset; e.g., Duñabeitia and Molinaro, 2014). The direct demonstration of orthographic pre-activation constitutes a theoretical advance of the present study, as previous evidence for predictive processing at the orthographic level had only been inferred indirectly from predictability effects after word onset (e.g., DeLong et al., 2019). Also, the findings described can be better explained by a predictive mechanism compared to spill-over effect.

In line with previous observations (Dikker and Pylkkänen, 2013; Fruchter et al., 2015; Wang et al., 2017), similar brain regions were activated during stimulus processing and pre-activation. In detail, source localization revealed lexical-semantic effects in frontal regions, replicating Fiebach et al. (2002) and Binder et al. (2003). For lexicality effects, this included the inferior frontal gyri as well as the temporal poles (previously implicated in lexical-semantic facilitation; Lau et al., 2013, 2016), which was expected given these regions form essential parts of the semantic network (e.g., Binder et al., 2009; Lambon Ralph et al., 2017). Furthermore, lexicality effects were localized to the cerebellum, which supports previous evidence that this brain structure is implicated in predictive processing of visual words (see review by Pleger and Timmann, 2018). In detail, it has been proposed that the cerebellum might serve a similar function during language processing as during motor control, i.e., representing internal models of contextually relevant stimuli such as words or objects (Moberget et al., 2014). Localizations of word frequency effects, besides frontal regions, also involved parietal regions (cf. Desai et al., 2018). Orthographic effects were localized to frontal (in the case of words) and parietal regions (in the case of pseudowords). Previous fMRI studies localized orthographic effects based on neighborhood measures (similar to the OLD20 measure used here) to regions implicated in orthographic processing (Yarkoni et al, 2008b; Braun et al., 2015) and/or to domain general regions (Fiebach et al., 2007; Yarkoni et al, 2008b; Braun et al., 2015). The sources found here fall into the latter category of regions previously associated with executive functions (e.g., Duncan, 2013), which might support (predictive) language processing (e.g., Ye and Zhou, 2009; Federmeier et al., 2020; Ryskin et al., 2020).

### 4.3 Implications for Models of Predictive Coding

The activation of linguistic representations before an expected linguistic stimulus is in line with the assumption of active prediction (as specified in predictive coding theory, e.g., Rao and Ballard, 1999). Kuperberg and Jaeger (2016) recently proposed that language comprehension involves predictive processes at multiple levels of the linguistic hierarchy. The present data support this hypothesis for orthographic and lexical-semantic, but not visual and phonological levels of word processing. Neither language-specific (Kuperberg and Jaeger, 2016) nor general theories of predictive coding (Friston, 2005) have discussed restrictions of predictive processing to certain levels of representation to the best of our knowledge. Therefore, this possibility should be explored further in future studies (see Gagl et al., 2020, for a recent example from pre-lexical orthographic processing).

Our data also indicate more temporally extended effects of orthographic pre-activation for words than pseudowords. This finding suggests that the presence of prior stimulus knowledge modulates predictive pre-activation. For example, it may be easier to activate (parts of) pre-existing mental representations in a top-down guided manner, rather than activate a newly assembled set of orthographic features of a previously unknown pseudoword. This notion is also compatible with the absence of sustained orthographic effects for pseudowords in the second half of the delay. The restriction of orthographic delay effects for pseudowords to very early time windows could reflect residual activation (spill-over) from processing the prime as opposed to pre-activation of orthographic target characteristics. On the other hand, this is not directly compatible with the observed change in the direction of orthographic effects between prime and delay (as discussed above). In this context, it is also interesting to note that pre-activation effects for lexical-semantic representations in the delay seem to temporally coincide with orthographic pre-activation for words but not pseudowords.

Besides predictive pre-activation, effects in the delay period might alternatively be interpreted as reflecting a working memory representation of the prime. This alternative explanation, in fact, cannot be excluded for the majority of studies investigating predictive pre-activation. As predictions unequivocally require memory (e.g., Bar, 2009), pre-activation cannot easily be differentiated from a mnemonic representation. For example, it has been suggested that working memory, just like prediction, might correspond to the activation of one’s ‘internal model of the world’ (e.g., Kok and de Lange, 2015). In addition, previous evidence indicated that individuals with higher working memory capacity show predictive behavior to a more substantial extent than persons with relatively lower working memory capacity, both in language processing (e.g., Huettig and Janse, 2016) and other domains (e.g., Cashdollar et al., 2017). Nevertheless, our observation that effects in the delay period were predominantly anti-correlated to effects elicited during prime processing rules out that these effects reflect a working memory representation of the prime (cf. Lee and Baker, 2016; Christophel et al., 2017). Instead, the positive correlation between the delay and target effects strengthens the interpretation that target characteristics are pre-activated during the delay.

Predictive coding theories assume that internal models can ‘explain away’ the known part of expected sensory inputs and that this part of a stimulus as a consequence does not have to be processed. Accordingly, brain responses elicited during perception should carry more information about the stimulus when perceiving unpredictable in contrast to predictable stimuli (Kok et al., 2012; Blank and Davis, 2016). We found this pattern for OLD20, where more significant effects were observed on the prime than the target. For lexical-semantic metrics, we observed more significant effects on the target than on the prime. This finding may be more compatible with ‘sharpening’, an alternative account of repetition priming effects according to which noisy parts of the signal are suppressed when sensory input is predictable, while the signal’s informative components are sharpened (e.g., Kok et al., 2012). The sharpening theory does not rule out predictive processing per se but represents an alternative mechanism of how predictive processing may be implemented (Kok et al., 2012; Walsh et al., 2020). Interestingly, previous studies in the domain of auditory word recognition found evidence for predictive coding as opposed to sharpening at the acoustic/phonological level (Blank and Davis, 2016; Sohoglu and Davis, 2020). These findings correspond well with the present evidence for predictive coding at a similarly early level – orthographic processing – in visual word recognition. In contrast, the observation that lexical-semantic context effects are rather in line with sharpening is a novel finding (note that the lexical-semantic level was not included in the investigations of Blank and Davis, 2016, and Sohoglu and Davis, 2020). Potentially, this reflects the high relevance of lexical-semantic information in word recognition. As the goal of word recognition is meaning access, it seems plausible that lexical-semantic features are not explained away but rather processed further, while a sharpening of these features nevertheless allows facilitated processing when predictable (i.e., as evident from the observed stronger reduction of behavioral response times for predictable words vs. pseudowords).

Also, the nature of correlations between delay and prime as well as delay and target seems to pose a challenge for predictive coding models. Predictive coding (Friston, 2005) assumes that the prediction contains relevant information about a stimulus, which is then suppressed when processing the expected stimulus. Accordingly, predictive coding would propose a positive correlation between unpredictable prime and delay representations, which was not observed for the majority of the present effects. Rather, the strong and sustained positive correlations between delay-period pre-activation and orthographic and lexical-semantic effects during the predictable target’s processing indicate that similar representations are active before and during predictable stimulus processing. In contrast, representations during unpredictable prime processing were more variable (i.e., of positive and negative effect direction). Again, this pattern suggests a sharpening mechanism (e.g., Kok et al., 2012) that reduces the variability of neuronal responses for predictable compared to unpredictable contexts. Tentatively, we propose that initial prime processing might be rather noisy as the prime is unpredictable. However, once the prime stimulus has been identified, readers are able to pre-activate a noise-free (‘sharpened’) representation of the upcoming expected target stimulus. This representation is also activated during the processing of the predictable target, explaining the observed correlation between delay and target representations.

### 4.4 Potential Limitations

Three methodological considerations may influence the interpretation of our findings: First, the fixed stimulus length (five letters) and resulting restriction of variance in the phonological predictor (number of syllables) may have limited our ability to observe behavioral effects related to visual and phonological processing levels. An unbiased investigation of potential contributions of these processing levels in future work may require more variable stimulus materials. Second, all results of the present study depend on the adequacy of the predictor variables for describing the representations at different processing levels. However, all stimulus measures used here are very well-established in psycholinguistics, and thus appear to be the best possible choices. Finally, the long delay interval between prime and target strongly diverges from fast-paced natural reading. We considered this long interval necessary in order to separate possible effects of pre-activation from stimulus processing; however, future research should aim at investigating context effects across representational levels in visual word recognition using a faster pacing of stimulus presentation or using natural reading combined with new analysis procedures (Dimigen and Ehinger, 2021).

### 4.5 Conclusion

We used linear mixed model analyses to show that in behavior and brain activation, context-dependent facilitation of visual word recognition relies most strongly on orthographic and lexical-semantic processes. At these two levels of linguistic representation, information is represented consistently and in a temporally coordinated manner in brain signals before the stimulus is presented. Our findings indicate that neuronal language processing systems actively prepare for the upcoming word and thereby increase word recognition efficiency via a sharpening mechanism.

## Author Notes

The research leading to these results has received funding from the European Community’s Seventh Framework Programme (FP7/2013) under grant agreement n° 617891 awarded to C.J.F. and a Marie Curie Fellowship under grant agreement n° 707932 awarded to B.G. We thank Isabelle Ehrlich, Anne Hoffmann, Jan Jürges, Rebekka Tenderra, and Stefanie Wu for their help with data acquisition, as well as two anonymous reviewers for helpful comments on a previous version of the manuscript. The authors declare no competing financial interests. Author contributions: S.E., B.G., and C.J.F. designed research; S.E. and B.G. performed research; S.E. analyzed data with contributions from B.G.; S.E., B.G., and C.J.F. wrote the paper.

## Supporting Information

### Supporting Materials. Correlations between stimulus parameters

**Figure S1.**
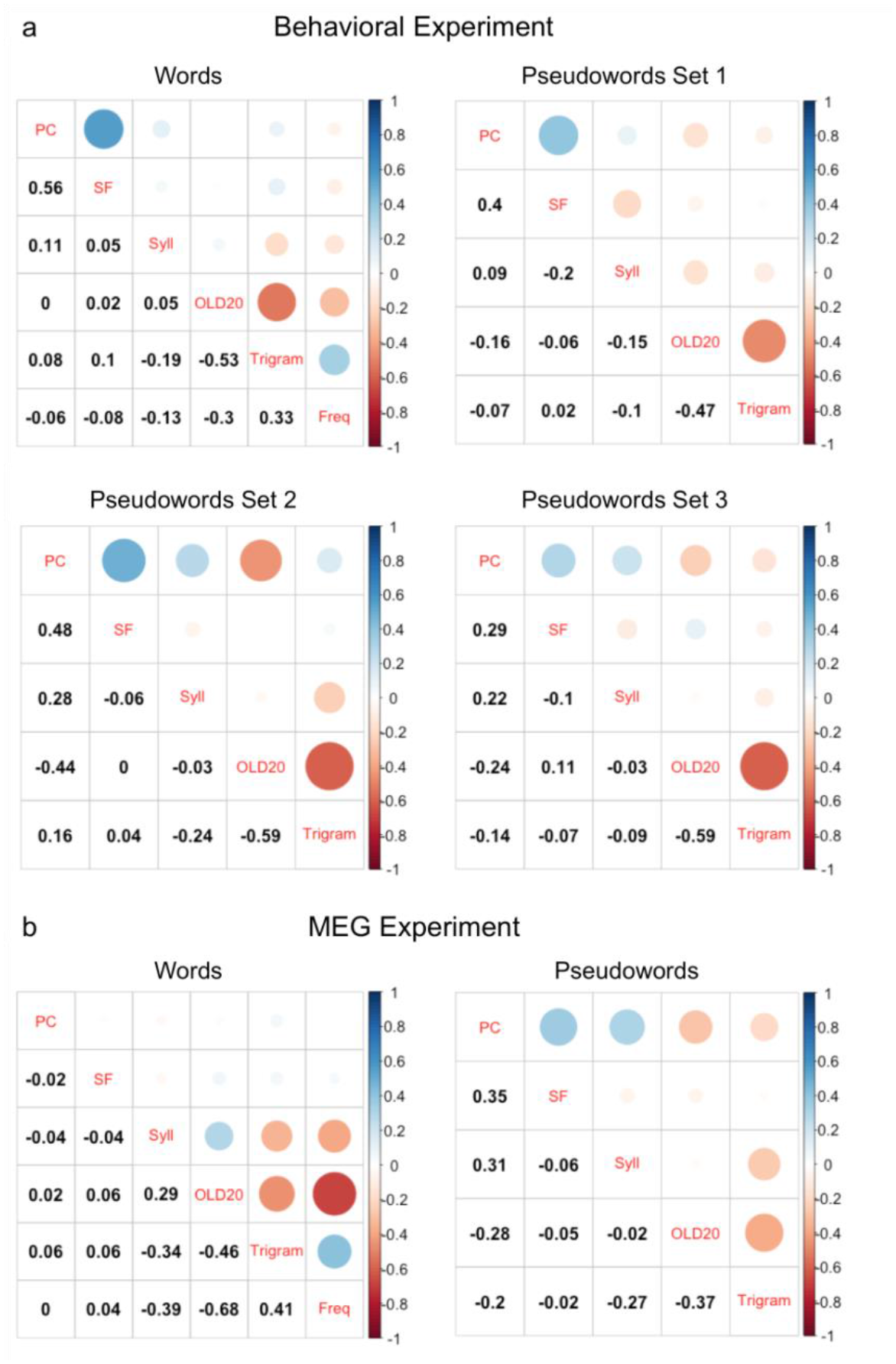
Correlations between stimulus parameters for words and pseudowords of (a) the behavioral and (b) the MEG experiment. PC: Perimetric complexity. SF: Number of simple features. Syll: Number of syllables. OLD20: Orthographic Levenshtein Distance 20. Trigram: Logarithmic trigram frequency. Freq: Logarithmic word frequency.

#### Supporting Results 1. Additional linear mixed modeling results for behavioral data

**Table S1.** Results of the full linear mixed model (LMM) of behavioral response times including – besides word frequency and OLD20 – also interactions of context (repeated target vs. isolated presentation) and lexicality with perimetric complexity, the number of simple features, the number of syllables, and trigram frequency

|  | *FE* | *SE* | *t* | *p* |
| --- | --- | --- | --- | --- |
| Context | **-0.275** | **0.013** | **-20.618** | **< 2e-16** |
| Word frequency | **-0.049** | **0.015** | **-3.271** | **0.00128** |
| Lexicality | -0.014 | 0.028 | -0.503 | 0.616 |
| OLD20 | -0.006 | 0.008 | -0.801 | 0.424 |
| Perimetric complexity | 0.007 | 0.008 | 0.899 | 0.369 |
| Trigram frequency | 0.008 | 0.008 | 0.966 | 0.335 |
| Simple features | -0.004 | 0.006 | -0.601 | 0.548 |
| Syllables | -0.004 | 0.010 | -0.429 | 0.669 |
| Trial order | **-0.025** | **0.002** | **-10.081** | **< 2e-16** |
| Word frequency x Context | **0.043** | **0.013** | **3.411** | **0.00065** |
| Lexicality x Context | **-0.139** | **0.025** | **-5.539** | **3.13e-08** |
| OLD20 x Context | -0.014 | 0.009 | -1.538 | 0.124 |
| OLD20 x Lexicality | 0.002 | 0.013 | 0.117 | 0.907 |
| Perimetric complexity x Context | 0.012 | 0.009 | 1.345 | 0.179 |
| Perimetric complexity x Lexicality | 0.001 | 0.013 | 0.084 | 0.933 |
| Trigram frequency x Context | 0.001 | 0.009 | 0.146 | 0.884 |
| Trigram frequency x lexicality | -0.006 | 0.014 | -0.438 | 0.662 |
| Simple features x Context | -0.002 | 0.007 | -0.265 | 0.791 |
| Simple features x Lexicality | -0.007 | 0.014 | -0.508 | 0.612 |
| Syllables x Context | 0.000 | 0.011 | 0.042 | 0.967 |
| Syllables x Lexicality | 0.001 | 0.012 | 0.121 | 0.904 |
| OLD20 x Context x Lexicality | **0.036** | **0.013** | **2.760** | **0.00579** |
| Perimetric complexity x Context x Lexicality | -0.014 | 0.013 | -1.106 | 0.269 |
| Trigram frequency x Context x Lexicality | 0.005 | 0.013 | 0.371 | 0.711 |
| Simple features x Context x Lexicality | 0.018 | 0.013 | 1.384 | 0.167 |
| Syllables x Context x Lexicality | 0.009 | 0.013 | 0.660 | 0.509 |

Bold numerals indicate significant effects, i.e., $p < .05$ . *FE* = fixed effect estimates, *SE* = standard error. Word frequency for pseudowords was set to zero.

**Table S2.** Results of post hoc linear mixed models (LMMs) resolving the three-way interaction of context with OLD20 and lexicality, as well as assessing the ‘classical’ word frequency effect which is typically estimated on words only, by estimating the two-way interactions of context with OLD20 and word frequency separately for words and pseudowords

|  | *Words* |  |  |  | *Pseudowords* |  |  |  |
| --- | --- | --- | --- | --- | --- | --- | --- | --- |
|  | *FE* | *SE* | *t* | *p* | *FE* | *SE* | *t* | *p* |
| Context | **-0.375** | **0.007** | **-56.192** | **< 2e-16** | **-0.314** | **0.008** | **-40.853** | **< 2e-16** |
| Word frequency | **-0.031** | **0.008** | **-3.715** | **0.00042** | - | - | - | - |
| OLD20 | -0.006 | 0.008 | -0.74 | 0.462 | **-0.013** | **0.006** | **-2.004** | **0.046** |
| Trial order | **-0.02** | **0.003** | **-6.249** | **4.5e-10** | **-0.03** | **0.004** | **-7.925** | **2.84e-15** |
| Word frequency x<br>Context | **0.025** | **0.007** | **3.69** | **0.00023** | - | - | - | - |
| OLD20 x Context | **0.017** | **0.007** | **2.32** | **0.020** | **-0.018** | **0.008** | **-2.378** | **0.017** |

Bold numerals indicate significant effects, i.e., $p < .05$ . *FE* = fixed effect estimates, *SE* = standard error.

**Table S3.**
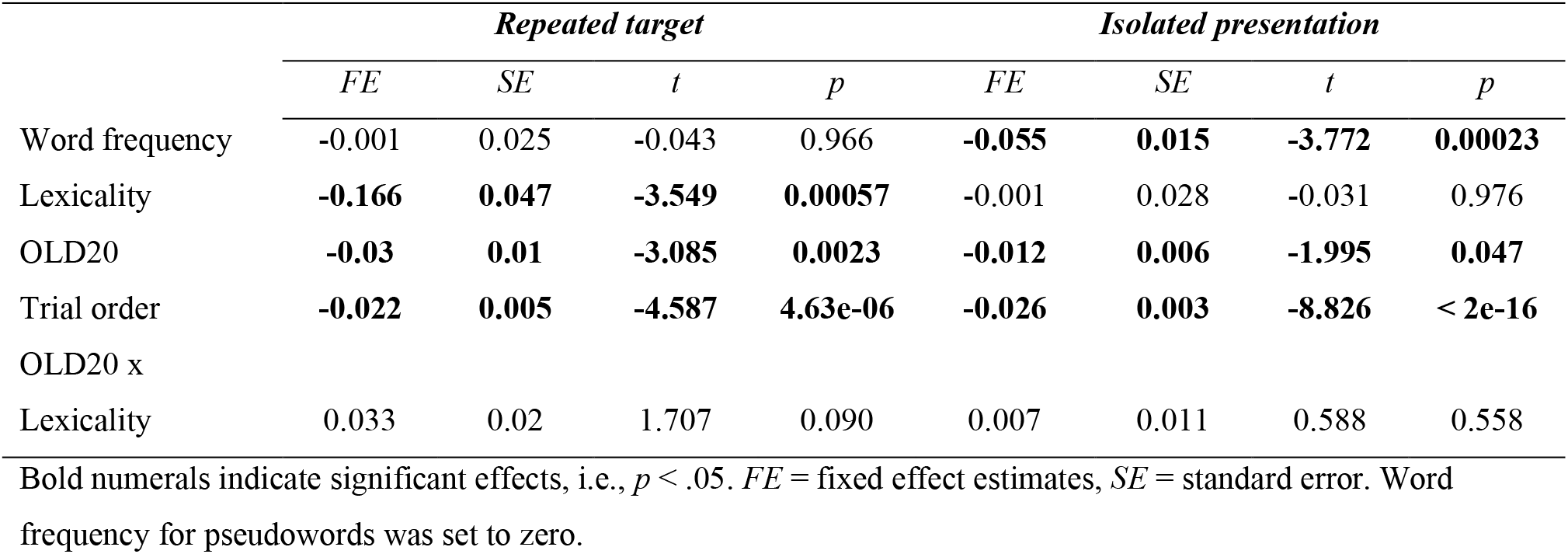
Results of post hoc linear mixed models (LMMs) resolving the interaction of context with word frequency as well as the three-way interaction of context with OLD20 and lexicality in behavioral response times by separately estimating word frequency effects and the OLD20 by lexicality interaction for repeated targets and isolated letter strings

#### Supporting Results 2. Pseudoword familiarization procedure and results

Our lexicality contrast mainly focused on the classical contrast between words and novel pseudowords, i.e., pseudowords that are unfamiliar to participants. However, these two groups do not only differ in the presence vs. absence of lexical-semantic information, but also in general familiarity, as only words are encountered in everyday life. To be able to dissociate between lexical-semantic and familiarity effects, we performed a pseudoword familiarization procedure. After familiarizing participants with a group of pseudowords, we assessed the lexicality contrast of words vs. familiar pseudowords. This contrast focuses on lexical-semantic differences between words and pseudowords as opposed to differences in general familiarity. Nevertheless, one has to acknowledge that familiar pseudowords might even be considered more familiar than words at the pre-lexical level, as both groups were initially matched on pre-lexical metrics (OLD20 and the number of syllables; cf. Eisenhauer et al., 2019) but only the familiar pseudowords were presented in the familiarization sessions. Still, we argue that effects that are found for words vs. novel pseudowords as well as for words vs. familiar pseudowords likely reflect differences in lexical-semantic processing. In the following, we first describe the familiarization procedure of the behavioral experiment and the MEG experiment, followed by the respective results.

##### Pseudoword familiarization procedure and results for the behavioral experiment

For the behavioral experiment, participants completed five pseudoword familiarization sessions over the course of three consecutive days. Pseudoword familiarization was adapted from Breitenstein et al. (2007) and followed the procedure for the behavioral experiment described in Eisenhauer et al. (2019). Each familiarization session consisted of reading aloud each to-be-familiarized pseudoword from a list (mean error rate across sessions: 0.7 %). Then, participants completed a paired-associate learning as well as a naming task. During paired-association, pseudowords were presented for 800 ms, followed by the presentation of an object image until button press or for a maximum duration of 1500 ms. At the beginning of each trial, the center of the screen was indicated as fixation point by the presentation of two vertical bars for 1000 ms. Each pseudoword was presented eight times, amounting to 960 trials in total. The ‘familiar pseudowords’ of interest for the present study were paired with a different object image (out of a set of 60 possible images) on each trial. The ‘semantic pseudowords’ not of interest for the present study were paired with the same object image in 75 % of trials. Participants were instructed to aim at learning a meaning for the pseudowords based on frequently co-occuring object images and were briefed that this would be possible only for half of the pseudowords. They were asked to respond to each object image via button press whether it matched the preceding pseudoword or not, stressing both response time and accuracy. On each trial, a red and a green bar presented on either side of the object image indicated which response hand should be used for a match or non-match response, respectively. At the start of the first session, participants practiced the task with at least ten trials until they confirmed they had understood the instructions. We observed that the average accuracy improved across sessions for all conditions. In detail, for matching semantic pseudowords and objects, accuracy increased from 44.1 % in session 1 to 91.9 % in session 5. For non-matching semantic pseudowords and objects, accuracy increased from 82.9 to 95.0 %. For familiar pseudowords, correct ‘non-match’ responses increased from 80.2 to 96.2 %.

In the naming task, each pseudoword was presented once and participants were instructed to name the associated object. In case they could not associate any object with a pseudoword, they were asked to respond ‘next’, which was the correct response for all familiar pseudowords. Naming accuracies improved from 22.6 % (range: 0 to 51.7 %) in session 1 to 80.9 % (range: 38.3 to 100 %) in session 5 for semantic, and from 85.0 % (range: 53.3 to 98.3 %) to 92.0 % (range: 45.0 to 100 %) for familiar pseudowords, confirming the majority of these pseudowords were not associated with a meaning.

At the end of the fifth familiarization session, participants completed an old/new recognition task in order to assess whether they recognized the trained pseudowords as familiar. In this task, each of the trained pseudowords was presented once until button press. In addition, a group of 120 filler pseudowords unfamiliar to the participants were presented. Participants pressed one of two buttons with their left or right index finger, indicating whether the presented pseudoword was familiar to them or not. Response hands were counterbalanced across participants. Due to a technical problem data of one participant were lost. Results confirmed that participants correctly recognized 93.9 % (range: 75.0 to 100 %) of semantic and 82.5 % (range: 21.7 to 100 %) of familiar pseudowords on average. In addition, 96.9 % (range: 90.0 to 100 %) of filler pseudowords were correctly classified as unfamiliar. All accuracies were significantly higher than chance level (50 %; one-sided t-test by item; semantic pseudowords: *t* = 79.259, df = 179, *p* < 2.2e-16; familiar pseudowords: *t* = 35.454, df = 179, *p* < 2.2e-16; filler pseudowords: *t* = 101.62, df = 119, *p* < 2.2e-16). The high accuracy values demonstrate that participants have reached a high level of familiarity with the familiar pseudowords. Following a break after the old/new recognition task, participants completed the repetition priming task on the same day.

##### Pseudoword familiarization procedure and results for the MEG experiment

Procedure and results for the pseudoword familiarization prior to the MEG experiment were described previously (Eisenhauer et al., 2019). Participants completed four familiarization sessions in the course of two consecutive days. In each session, participants read out pseudowords aloud from a list (mean error rate across sessions: 0.7 %) and performed an old/new recognition task as described above, presenting each of the 60 to be familiarized pseudowords two times, as well as 120 new filler pseudowords per session (no semantic pseudowords were presented). Due to a technical issue, data of sessions 3 and 4 were lost for one participant. Accuracies improved from 61.4 % in session 1 (range: 0.8 to 86.7 %) to 92.3 % in session 4 (range: 70.0 to 99.2 %) for familiar pseudowords, and from 77.8% (range: 16.7 to 99.2 %) to 91.3 % (range: 59.2 to 99.2 %) for filler pseudowords. Accuracies of the final session were significantly higher than chance for both conditions (50 %; one-sided t-test by item; familiar pseudowords: *t* = 51.4, df = 59, *p* < 2.2e-16; filler pseudowords: *t* = 34.7, df = 119, *p* < 2.2e-16).

#### Supporting Results 3. Linear mixed modeling results for the lexicality contrast of words vs. familiar pseudowords

##### Behavioral results

**Figure S2.**
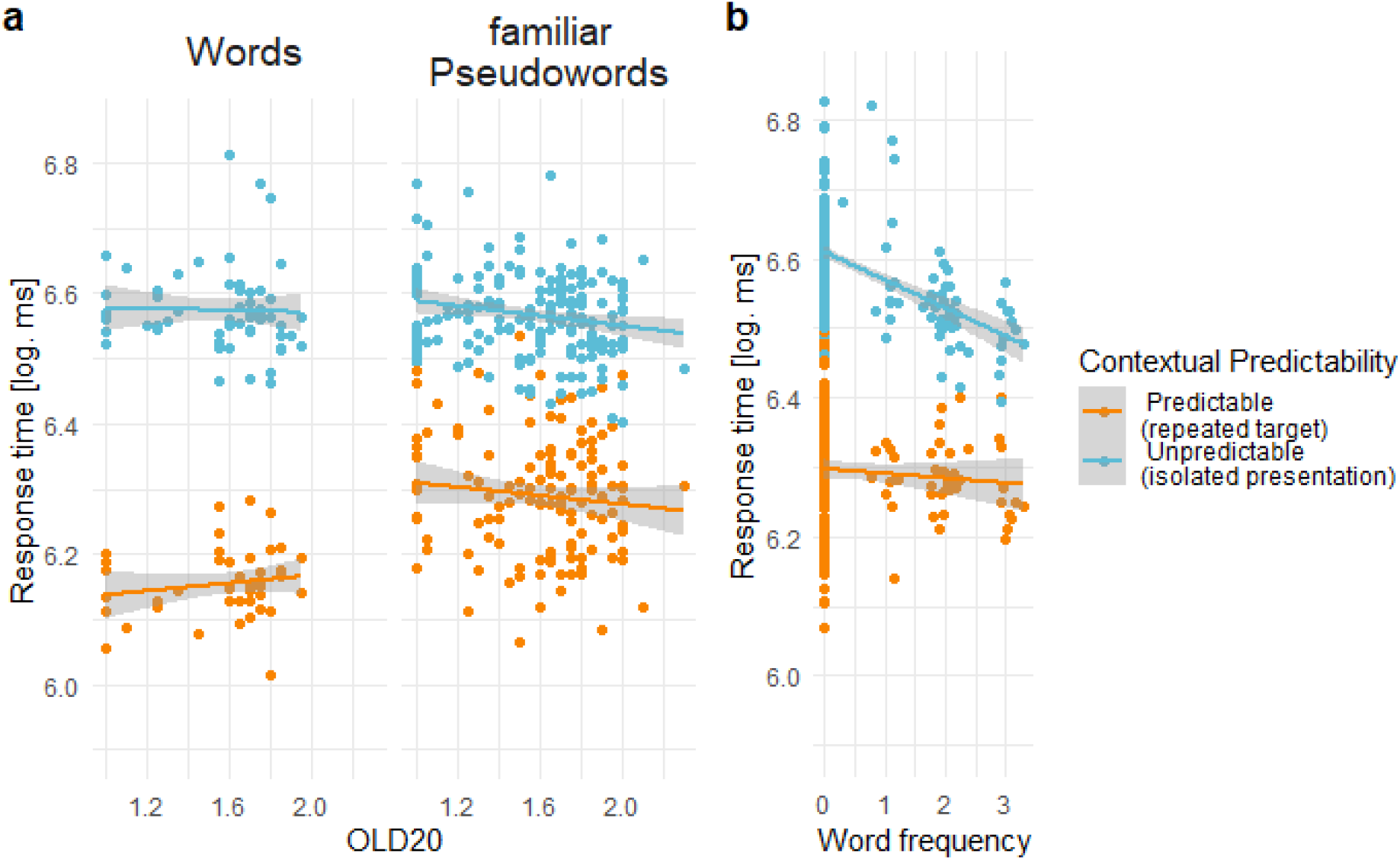
Behavioral results reflecting context-dependent facilitation for words and familiar pseudowords. Comparison of response times representing (a) orthographic (OLD20 effect separated for words and familiar pseudowords) and (b) lexical processing (word frequency effect) when stimuli were either predictable from context (i.e., repeated targets; orange lines and dots) or not (isolated presentation; blue lines and dots). Note that lower OLD20 values reflect higher word likeness, and, thus, orthographic familiarity. Displayed are logarithmic response times that represent the partial effects estimated from linear mixed models. Dots represent items averaged across participants.

**Table S4.** Results of the linear mixed model (LMM) investigating context effects (repeated target vs. isolated presentation) on word frequency, OLD20, and lexicality (words vs. familiar pseudowords) in behavioral response times

|  | *FE* | *SE* | *t* | *p* |
| --- | --- | --- | --- | --- |
| Context | **-0.273** | **0.013** | **-20.316** | **< 2e-16** |
| Word frequency | **-0.046** | **0.013** | **-3.550** | **0.0005** |
| Lexicality | -0.017 | 0.024 | -0.690 | 0.491 |
| OLD20 | -0.009 | 0.005 | -1.584 | 0.114 |
| Trial order | **-0.026** | **0.002** | **-10.027** | **< 2e-16** |
| Word frequency x Context | **0.039** | **0.012** | **3.220** | **0.001** |
| Lexicality x Context | **-0.135** | **0.024** | **-5.577** | **2.52e-08** |
| OLD20 x Context | -0.001 | 0.007 | -0.170 | 0.865 |
| OLD20 x Lexicality | 0.003 | 0.010 | 0.332 | 0.740 |
| OLD20 x Context x Lexicality | 0.017 | 0.011 | 1.498 | 0.134 |

Bold numerals indicate significant effects, i.e., $p < .05$ . *FE* = fixed effect estimates, *SE* = standard error. Word frequency for pseudowords was set to zero. Post hoc tests are shown in Tables S5 to S7.

**Table S5.** Results of post hoc linear mixed models (LMMs) estimating word frequency effects and the OLD20 by lexicality (words vs. familiar pseudowords) interaction on behavioral response times separately for repeated targets and isolated letter strings

|  | *Repeated target* |  |  |  | *Isolated presentation* |  |  |  |
| --- | --- | --- | --- | --- | --- | --- | --- | --- |
|  | *FE* | *SE* | *t* | *p* | *FE* | *SE* | *t* | *p* |
| Word frequency | -0.001 | 0.022 | -0.067 | 0.947 | **-0.055** | **0.014** | **-3.805** | **0.0002** |
| Lexicality | **-0.016** | **0.042** | **-3.914** | **0.0001** | 0.003 | 0.027 | 0.107 | 0.915 |
| OLD20 | -0.010 | 0.009 | -1.116 | 0.266 | **-0.011** | **0.006** | **-1.777** | **0.077** |
| Trial order | **-0.021** | **0.005** | **-4.178** | **3e-05** | **-0.032** | **0.003** | **-10.300** | **< 2e-16** |
| OLD20 x Lexicality | 0.013 | 0.018 | 0.734 | 0.465 | 0.006 | 0.011 | 0.491 | 0.624 |

Bold numerals indicate significant effects, i.e., $p < .05$ . *FE* = fixed effect estimates, *SE* = standard error. Word frequency for pseudowords was set to zero.

**Table S6.** Results of post hoc linear mixed models (LMMs) estimating the two-way interaction of context with OLD20 for familiar pseudowords in behavioral response times

|  | *Familiar Pseudowords* |  |  |  |
| --- | --- | --- | --- | --- |
|  | *FE* | *SE* | *t* | *p* |
| Context | **-0.308** | **0.008** | **-36.990** | **< 2e-16** |
| OLD20 | -0.009 | 0.006 | -1.490 | 0.137 |
| Trial order | **-0.033** | **0.004** | **-8.168** | **4.06e-16** |
| OLD20 x Context | -0.002 | -0.008 | -0.198 | 0.843 |

Bold numerals indicate significant effects, i.e., $p < .05$ . *FE* = fixed effect estimates, *SE* = standard error.

**Table S7.**
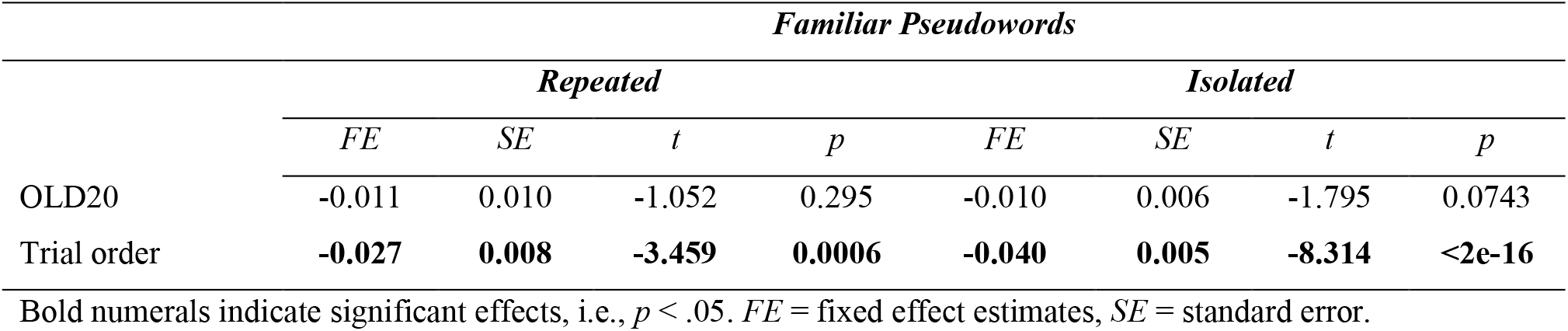
Results of the post hoc linear mixed model (LMM) investigating OLD20 effects on behavioral response times separately for repeated and isolated familiar pseudowords

##### MEG results

**Figure S3.**
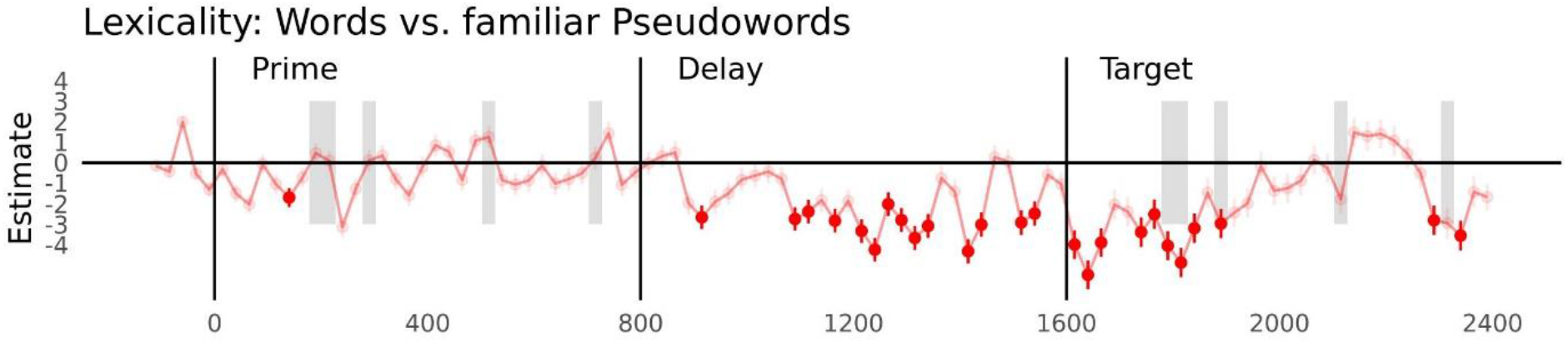
Lexicality effect reflecting the contrast between words and familiar pseudowords in MEG-measured brain responses. Displayed are the estimates of linear mixed models ± *SE* across time, including baseline, prime, delay, and target time windows. The salient red dots represent significant time points, excluding un-estimable models. Gray shading represents time points of a significant interaction between lexicality and context (prime vs. target). Note that the models were identical to the main model of words vs. novel pseudowords, i.e., the interaction of lexicality and context with OLD20 was also controlled for to allow comparability. However, OLD20 effects are not presented here as the interactions with OLD20 did not reach significance in the behavioral model of words and familiar pseudowords.

#### Supporting Results 4. Behavioral linear mixed modeling results comparing response times for repeated targets, isolated letter strings and non-repeated targets

**Figure S4.**
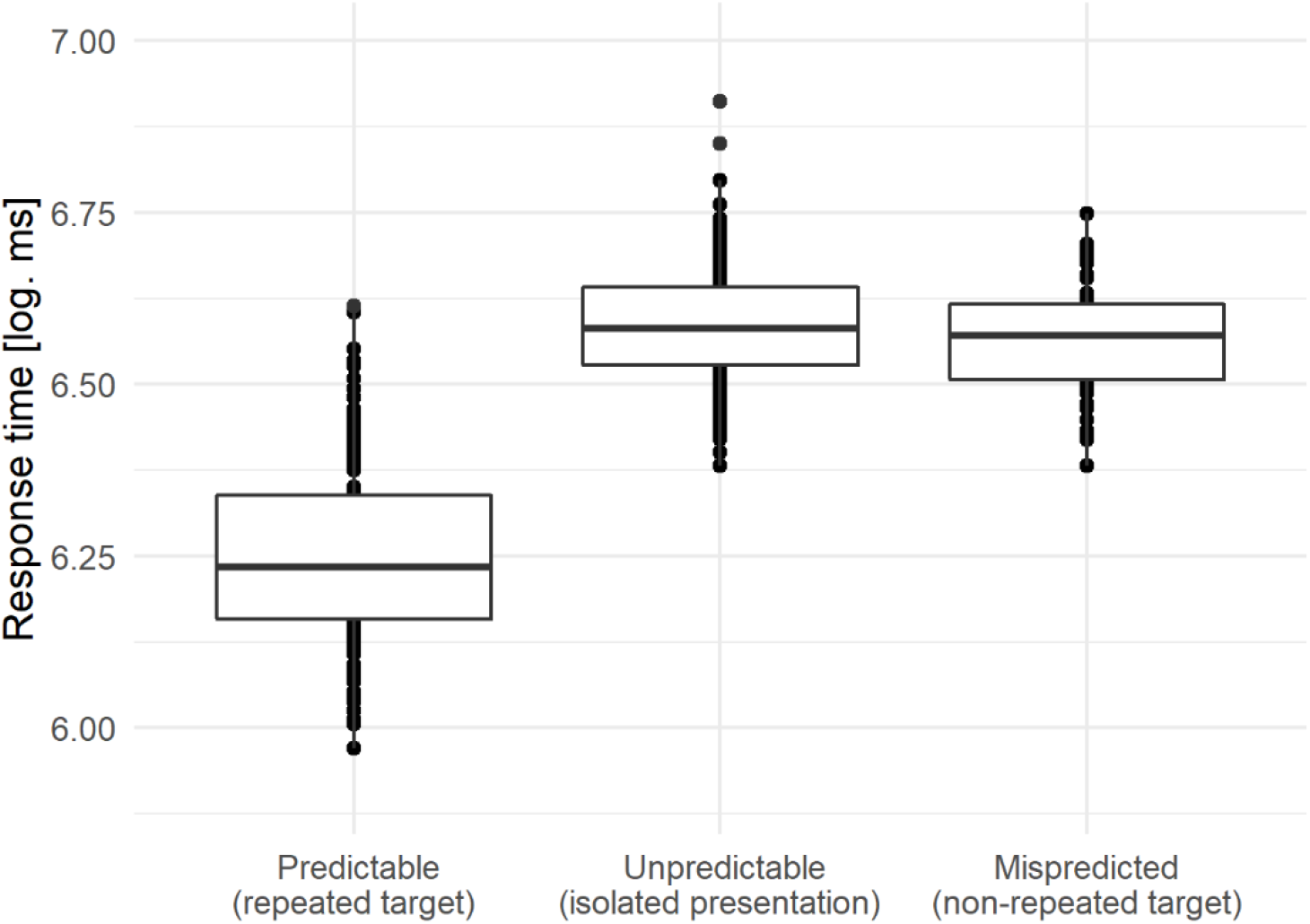
Behavioral logarithmic response times to predictable (repeated targets), unpredictable (isolated items) and mispredicted letter strings (non-repeated targets). Dots represent items averaged across participants.

**Table S8.**
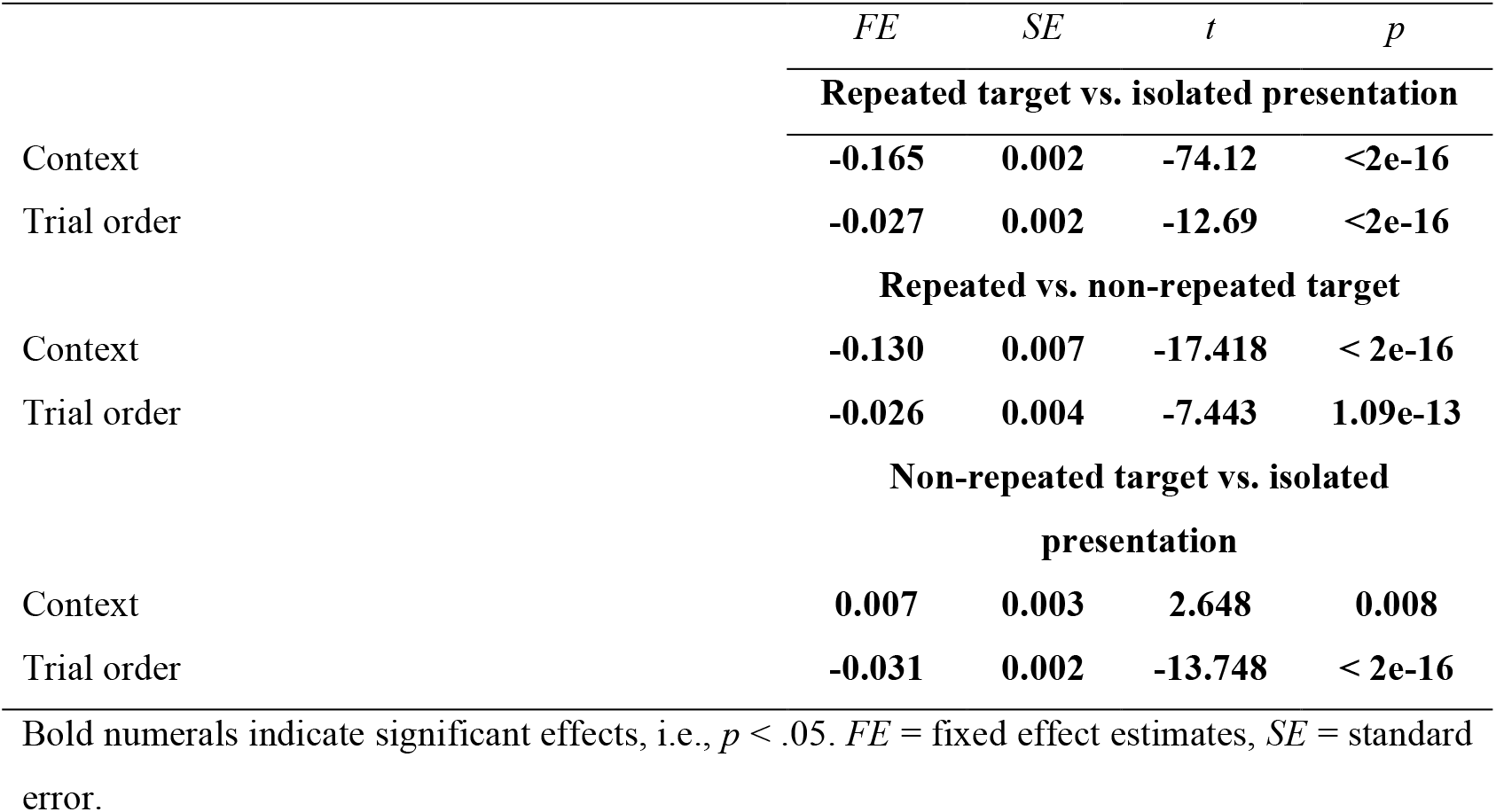
Results of the linear mixed model (LMM) investigating different context effects, i.e., repeated target vs. isolated presentation; repeated target vs. non-repeated target; and non-repeated target vs. isolated presentation in behavioral response times

|  | *FE* | *SE* | *t* | *p* |
| --- | --- | --- | --- | --- |
| **Repeated target vs. isolated presentation** |  |  |  |  |
| Context | **-0.165** | **0.002** | **-74.12** | **<2e-16** |
| Trial order | **-0.027** | **0.002** | **-12.69** | **<2e-16** |
| **Repeated vs. non-repeated target** |  |  |  |  |
| Context | **-0.130** | **0.007** | **-17.418** | **< 2e-16** |
| Trial order | **-0.026** | **0.004** | **-7.443** | **1.09e-13** |
| **Non-repeated target vs. isolated presentation** |  |  |  |  |
| Context | **0.007** | **0.003** | **2.648** | **0.008** |
| Trial order | **-0.031** | **0.002** | **-13.748** | **< 2e-16** |

Bold numerals indicate significant effects, i.e., $p < .05$ . *FE* = fixed effect estimates, *SE* = standard error.

#### Supporting Results 5. Multivariate pattern decoding analysis and results

We had initially favored multivariate pattern analysis (MVPA) as method of choice for the investigation of pre-activation effects in the MEG data, as previous studies had successfully decoded linguistic information from M/EEG data even in the absence of external stimulation (Simanova et al., 2015; Heikel et al., 2018). However, with this procedure we observed that the statistical power was limited and only resulted in significant findings for the strong and categorical lexicality effect during prime processing (Fig. S5). The absence of decodability of OLD20 and word frequency, i.e., the two investigated parametric variables, from MEG responses was unexpected (i) as both effects were found reliable in our behavioral data, and (ii) given that word frequency and orthographic similarity have in previous work been found to modulate M/EEG responses to words and pseudowords (e.g., Embick et al., 2001; Assadollahi and Pulvermüller, 2003; Vergara-Martínez and Swaab, 2012; Dufau et al., 2015; Gagl et al., 2016; Carrasco-Ortiz et al., 2017). Effects of OLD20 and word frequency should thus at least be found to reliably modulate brain activity during the presentation of the prime. As this was not the case, we reasoned that the MVPA procedure might either lack the statistical power to capture these effects, or that confounding effects (e.g., due to correlated word characteristics) may have obscured the effects of interest. To overcome these potential limitations, we investigated effects of the three stimulus characteristics of interest on MEG-measured brain activation using linear mixed models (LMMs), analogous to the statistical approach used for analyzing the data from the behavioral experiment. To compare the two analysis approaches, a power analysis was calculated (Fig. S6).

MVPA was performed using the SlidingEstimator function of the open-source package MNE-Python (https://martinos.org/mne/stable/index.html; Gramfort et al., 2014). Analyses involving the target time window were conducted exclusively on repetition trials (i.e., trials with identical primes and targets), whereas analyses of baseline, prime, and delay intervals included all available data (i.e., non-repetition and repetition trials). Decoding analyses were performed for each participant on the z-transformed multivariate pattern of neuronal responses measured at each of the 269 MEG sensors, which served as features. We used linear decoders (cf. King et al., 2020) with default parameters (alpha = 1) implemented in the Python-based scikit-learn package (Pedregosa et al., 2011). For the binary lexicality contrast (i.e., words vs. pseudowords), a Logistic Regression decoder was used on data stratified to contain 50 % of trials per condition, while continuous parameters were predicted using a Ridge Regression decoder (Tikhonov et al., 1977). In a 5-fold cross-validation approach, the decoder was trained, in each iteration, on 80 % of the data and then used to predict the remaining 20 % of unseen data, such that each trial was used for testing only once. In detail, trials were randomly partitioned into five folds (with the constraint for the lexicality contrast that training and testing data both contained 50 % of trials from each letter string condition), each of which served as testing data once after the decoder had been trained on data from the remaining four folds, thereby allowing us to derive a prediction for all trials in the dataset. To counteract a potential influence of the assignment of trials into folds, this procedure was repeated 100 times with trials randomly assigned to the five partitions of data.

**Figure S5.**
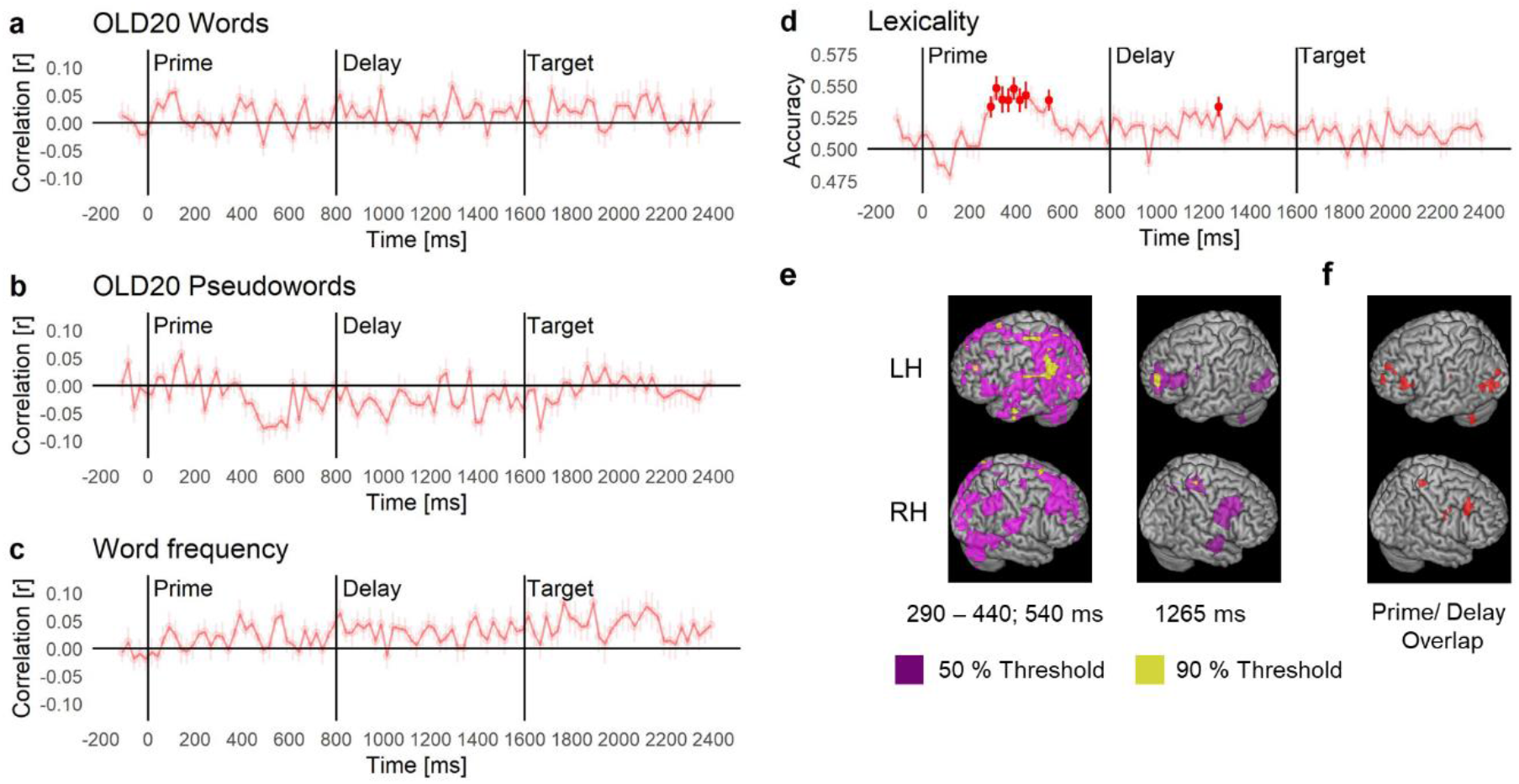
Multivariate pattern analysis (MVPA) decoding performance quantified as the Pearson correlation coefficient between actual and predicted parameter values from ridge regression decoders ± SEM (a-b) or the area under the receiver operating characteristic curve from logistic regression decoders (d) for (a) OLD20 in words, (b) OLD20 in pseudowords, (c) word frequency (in words only), and (d) lexicality. Salient dots represent significant time points (Bonferroni-corrected one-sided Wilcoxon signed-rank test against chance, i.e., a correlation coefficient of zero or an accuracy of .5). The difference between prime and target did not reach significance for any contrast or time point (Bonferroni-corrected). (e) Source topographies display event-related field based sources of significant time points of the lexicality contrast, thresholded at 50 % of the overall maximal activation (violet) within the left (LH) and right (RH) hemisphere. Additionally, source activations thresholded at 90 % of the individual peak per time point are shown (yellow). For the prime contrast, source activations were averaged across significant time points. Note that no control of confounding factors, as for the LMM-based source analysis, was possible. (f) Overlapping sources during prime and delay time windows across all significant time points, thresholded at 50 % of the overall peak. See Table S9 for source coordinates.

**Table S9.** MNI coordinates and source location of the strongest clusters as well as the overlapping clusters for the lexicality contrast presented in Figure S4

| Time [ms] | Cluster | Peak MNI coordinates |  |  | Region of cluster |
| --- | --- | --- | --- | --- | --- |
|  |  | x | y | z |  |
| Prime |  |  |  |  |  |
| 290 to 440;<br>540 | 1 | -53.5 | -52.5 | 38.5 | left angular gyrus/ inferior parietal lobule |
|  | 2 | -30.5 | -32.5 | 68.5 | left postcentral gyrus |
|  | 3 | 2.5 | 11.5 | 68.5 | right superior frontal gyrus |
| Delay |  |  |  |  |  |
| 1265 | 1 | -44.5 | 50.5 | 19.5 | left frontal pole |
|  | 2 | 30.5 | -30.5 | 69.5 | right postcentral gyrus |
| Prime/Delay Overlap |  |  |  |  |  |
|  | 1 | -39.5 | -80.5 | 4.5 | left lateral inferior occipital cortex |
|  | 2 | 44.5 | 20.5 | 40.5 | right middle frontal gyrus |
|  | 3 | -51.5 | 29.5 | 14.5 | left inferior frontal gyrus pars triangularis |

Prime and delay clusters are based on the source activation thresholded at 90 % of the individual peak. Overlap estimation included all significant sources of the respective time windows, thresholded at 50 % of the overall peak.

#### Supporting Results 6. Power comparison between linear mixed modeling and MVPA decoding analysis

To estimate the statistical power for LMM and MVPA analyses, we implemented a Monte-Carlo simulation-based power analysis in R: As a first step, data (behavioral as well as MEG, independent analyses) were randomly sampled from the original datasets (the baseline in case of the MEG data), and split into two halves respecting the random effect structure of the dataset (e.g., the number of participants, and, in case of the LMM-based analysis, sensors). For the LMM-based analysis, we then added to one half of the dataset a specific value reflecting the to-be-tested effect size and conducted statistical analyses as for the original dataset (including Bonferroni correction). For the MVPA analysis, we sampled from the classification accuracies of lexicality, added the critical effect size value, and then tested statistically against .5, i.e., the chance level for this analysis. We re-estimated these analyses 10,000 times for each effect size level (i.e., d values from 0.1 to 1), dataset (behavioral and MEG), and statistical analysis (LMM and MVPA), and recorded the p-values and estimated effect sizes from all tests. The probability of a test being significant at any given effect size represents the power. The mean of all effect size estimates from the significant tests allowed us to estimate the effect size bias, i.e., the likelihood that a significant finding represents an overestimated effect size. Inspection of the power curves (Fig. S6a) reveals that for the LMM analysis of the behavioral data, the present study achieved a power of greater than .9 at small effect sizes of .3. Thus, the likelihood of obtaining a significant result if a small effect exists in the population was greater than 90 %. This clearly reflects the benefit of modeling data at the level of the individual trial (as opposed to the ‘classical’ approach of aggregating trials by subject and condition before calculating statistics; see also Baayen et al., 2008). For the same effect size, the LMM analysis of brain data resulted in a power of around .6 and, alarmingly, for the MVPA analysis, the power of obtaining a statistically significant result was lower than .1. This finding confirms our intuition after the inspection of our brain activation results, i.e., that the MVPA analysis is much less powered. The consequences are severe, because there is a substantial possibility that significant MVPA findings may vastly overestimate the size of the true effect (Fig. S6b). This phenomenon is a frequent but unwanted effect of the correct practice of neglecting non-significant tests, so that significant effects of an underpowered study are often over-estimated in their effect size (e.g., Yarkoni, 2009). LMM analysis is much less susceptible to such overestimations, mainly because it can explicitly estimate crossed random effects (such as subjects and items) by modeling all data at the same time, while MVPA (and other methods) requires aggregation of data points (for more detailed discussions, see Baayen et al., 2008; Bates et al., 2018). The higher power and reduced bias of the behavioral compared to the MEG analyses further supports our procedure of constraining MEG investigations based on behavioral results; this reduces the possibility of false-positive effects, which is generally high in very high-dimensional datasets.

**Figure S6.**
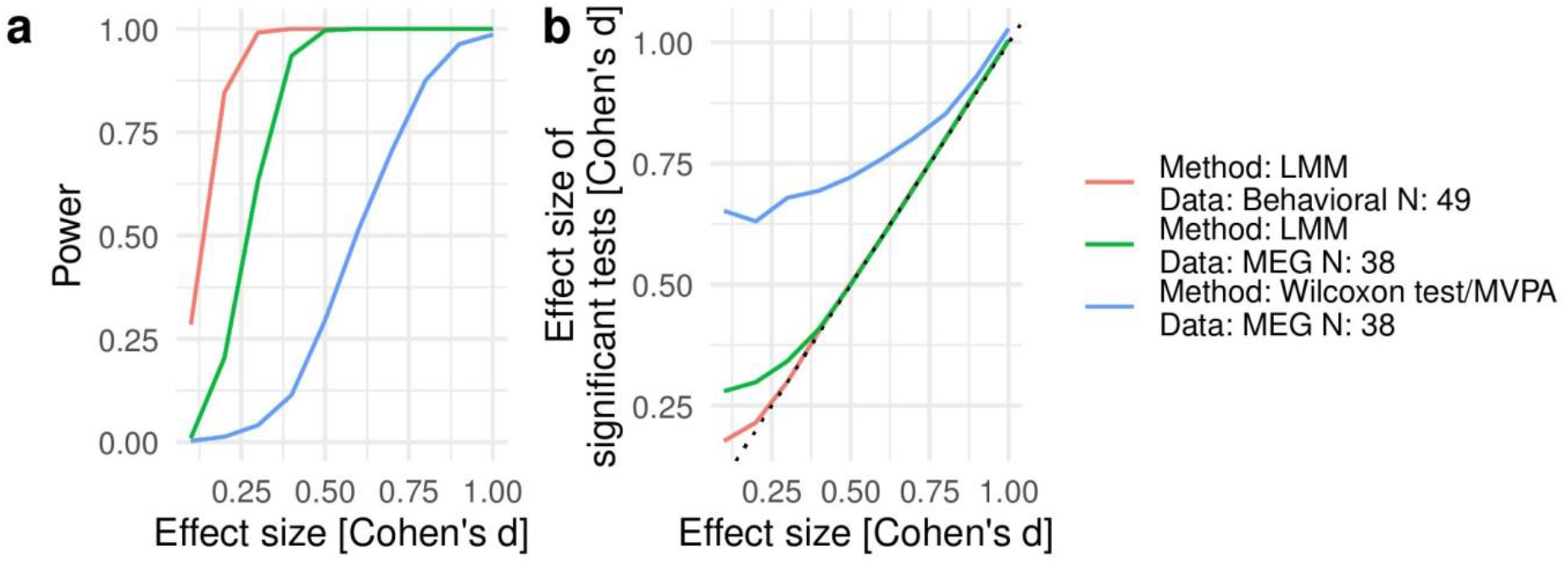
(a) Power curve and (b) effect size inflation for the behavioral analysis and LMM as well as MVPA procedures used to analyze the MEG data.

#### Supporting Results 7. MNI coordinates and source locations of sources presented in Figure 4

**Table S10.**
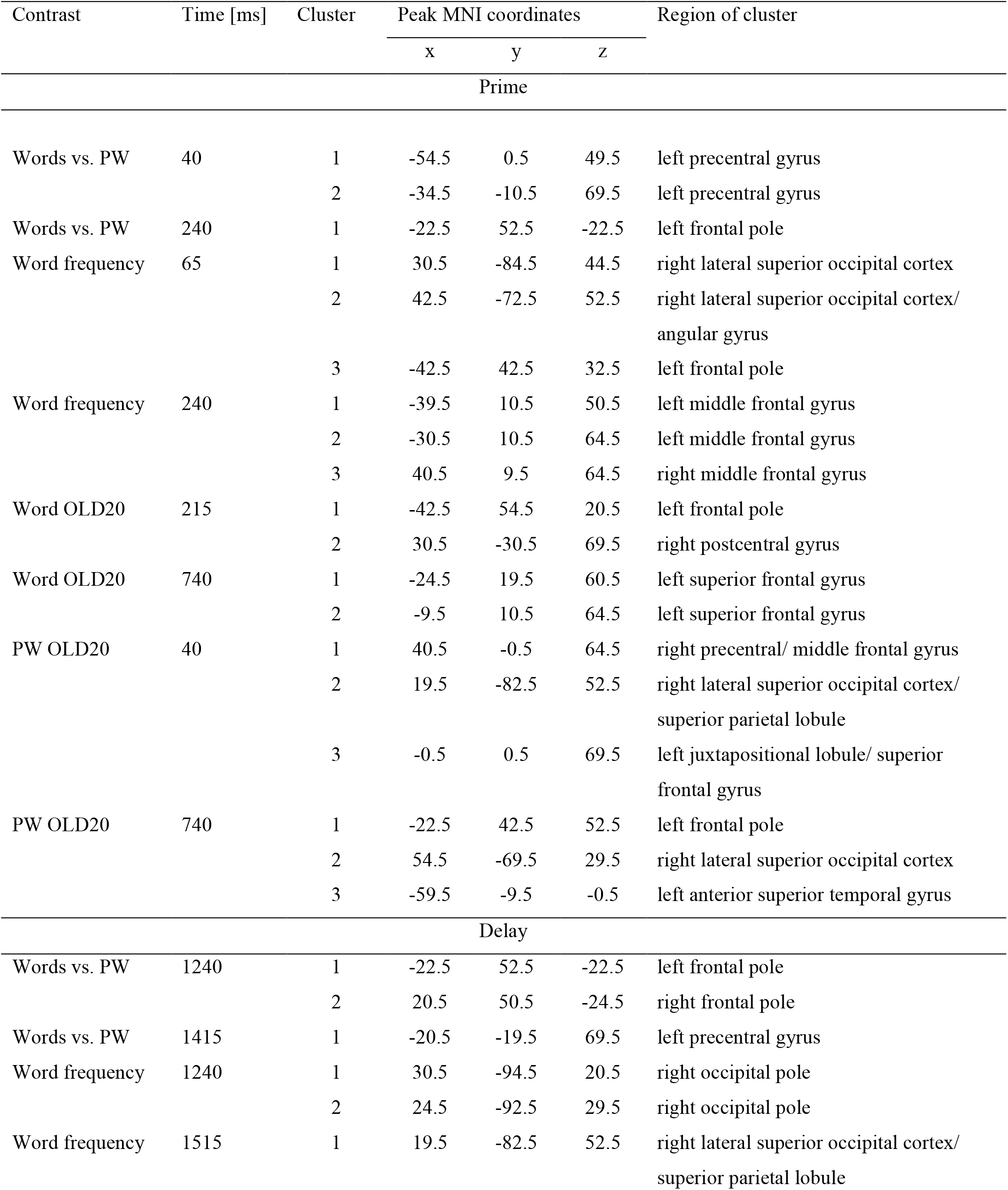
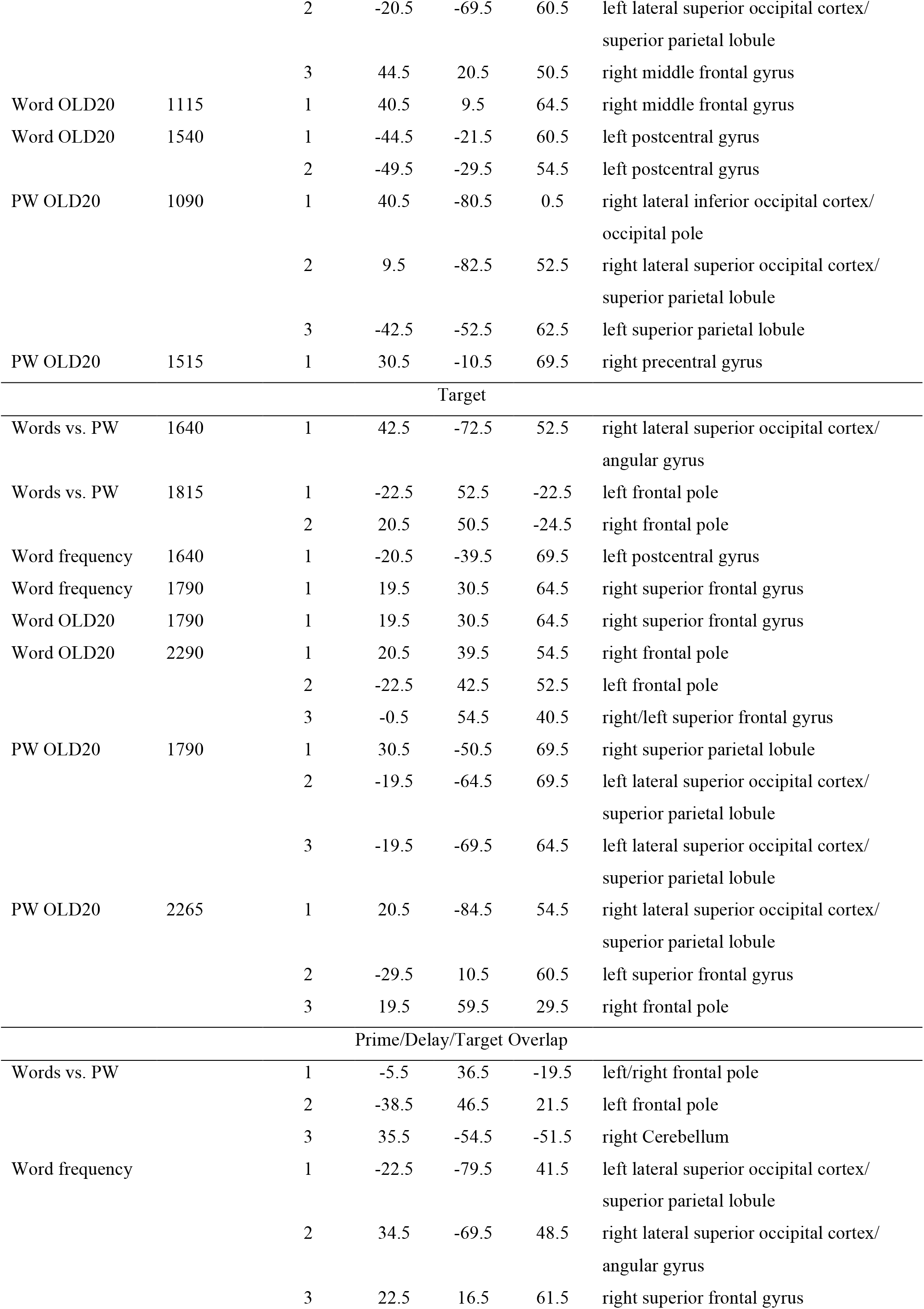
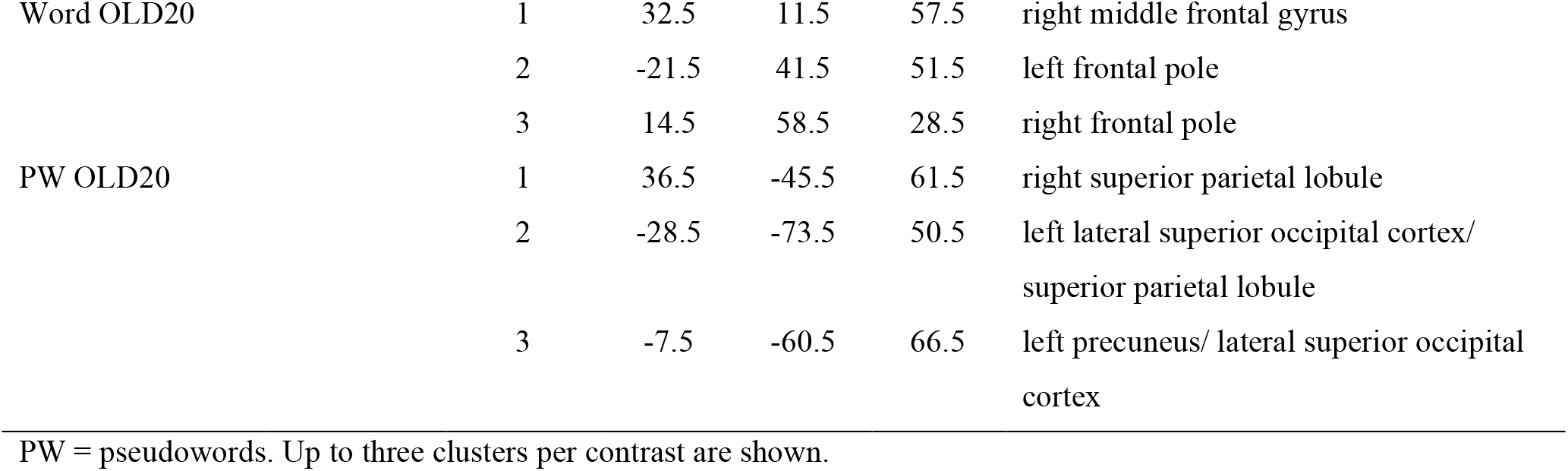
MNI coordinates and source locations of the strongest clusters (thresholded at 90 % of the individual peak) presented in Figure 4, as well as of the largest overlapping clusters (based on both a 90 % threshold of the individual peak as well as a 50 % threshold of the overall peak) across prime, delay and target

